# MAT2a and AHCY inhibition disrupts antioxidant metabolism and reduces glioblastoma cell survival

**DOI:** 10.1101/2024.11.23.624981

**Authors:** Emma C. Rowland, Matthew D’Antuono, Anna Jermakowicz, Nagi G. Ayad

**Affiliations:** Georgetown University, Lombardi Comprehensive Cancer Center, 3970 Reservoir Rd NW Washington D.C. 20007, United States of America

**Keywords:** methionine, metabolism, glioblastoma, oxidative stress, mitochondria, lipid peroxidation, metabolomics

## Abstract

Glioblastoma (GBM) is a highly aggressive primary malignant adult brain tumor that inevitably recurs with a fatal prognosis. This is due in part to metabolic reprogramming that allows tumors to evade treatment. We therefore must uncover the pathways mediating these adaptations to develop novel and effective treatments. We searched for genes that are essential in GBM cells as measured by a whole-genome pan-cancer CRISPR screen available from DepMap and identified the methionine metabolism genes *MAT2A* and *AHCY*. We conducted genetic knockdown, evaluated mitochondrial respiration, and performed targeted metabolomics to study the function of these genes in GBM. We demonstrate that *MAT2A* or *AHCY* knockdown induces oxidative stress, hinders cellular respiration, and reduces the survival of GBM cells. Furthermore, selective MAT2a or AHCY inhibition reduces GBM cell viability, impairs oxidative metabolism, and changes the metabolic profile of these cells towards oxidative stress and cell death. Mechanistically, MAT2a or AHCY regulates spare respiratory capacity, the redox buffer cystathionine, lipid and amino acid metabolism, and prevents DNA damage in GBM cells. Our results point to the methionine metabolic pathway as a novel vulnerability point in GBM.

**Significance:** We demonstrated that methionine metabolism maintains antioxidant production to facilitate pro-tumorigenic ROS signaling and GBM tumor cell survival. Importantly, targeting this pathway in GBM can potentially reduce tumor growth and improve survival in patients.

## Introduction

Glioblastoma (GBM) is the most common malignant primary brain tumor in adults^1^. Despite efforts to identify new treatments for GBM, prognosis is dismal with a survival rate of approximately 15 months following diagnosis^2^. The standard of care involves tumor resection along with a combination of temozolomide (TMZ), radiation treatment, and tumor treating fields^3^. Unfortunately, recurrence is nearly inevitable even following maximum bulk tumor resection. Treatments are ineffective in part due to the metabolic reprogramming that occurs in GBM cells^4^. Studies suggest that GBM relies on oxidative metabolism (including oxidative phosphorylation and fatty acid oxidation) to a greater extent than glycolysis^5^, and that treatment insult may inadvertently promote a metabolic shift towards oxidative metabolism to facilitate tumor progression^6–9^. Therefore, there is a need to identify novel mechanisms or actionable metabolic targets in GBM to circumvent or exploit this dependency.

One critical pathway that has been identified in GBM tumorigenesis and progression is one-carbon metabolism^10^, which involves catabolism and recycling of the essential amino acid methionine to regulate several key processes including immunity, lipid metabolism, and antioxidant production^11^. It was discovered that GBM tumors are largely dependent on methionine metabolism through tumor imaging using positron emission tomography, which revealed considerable ^11^C-methionine uptake compared to normal brain tissue^12^. This finding has been supported by reduced GBM growth through methionine media depletion *in vitro* and dietary methionine restriction *in vivo*^13–15^. Additionally, methionine restriction has been shown to improve the efficacy of TMZ in an orthotopic nude mouse model^16^, however the mechanism by which this occurs and the process through which methionine promotes GBM tumor progression is not well understood.

Here, we identify two enzymes involved in methionine metabolism, MAT2a and AHCY, which are found to be essential for GBM growth and redox balance. No known studies to date have interrogated these two enzymes and their involvement in GBM antioxidant metabolism. We demonstrate that both MAT2a and AHCY are indispensable for proper mitochondrial function in GBM and are necessary to protect cancer cells against oxidative stress. Thus, our study designates these two metabolic enzymes as important candidates for targeted therapy in GBM.

## Results

### Genes encoding for MAT2a and AHCY are essential in GBM and other CNS tumors

We sought to identify whether specific methionine cycle-related metabolic genes are essential in GBM cell growth and survival. To explore this, we probed the Cancer Dependency Map or DepMap (https://depmap.org/portal)^17^. DepMap is an online public database provided by the Broad Institute that utilizes large-scale functional genomics profiling in thousands of cancer cell lineages to elucidate gene essentiality. Their findings are based on pooled whole genome CRISPR-Cas9 screens across multiple cell lineages, primary diseases, and disease subtypes. A gene is deemed essential based on a calculated dependency score: an effect score of less than zero indicates reduced cell growth while an effect score of -0.5 or lower indicates induced cell death upon gene knockout. We investigated fifteen primary metabolic genes involved in the methionine pathway in the DepMap database (**Figures 1a-c**). From this analysis, we identified two genes, *MAT2A* and *AHCY*, that upon genetic knockout were associated with reduced cell growth for diffuse glioma (*MAT2A* gene effect -1.4, p= 3.4×10^-13^; *AHCY* gene effect -.335, p= 3.6×10^-9^) and GBM (*MAT2A* gene effect -1.38, p= 1.7×10^-10^; *AHCY* gene effect -.305, p= 2.7×10^-8^) cell lineages (**Figures 1d, e**). *MAT2A* encodes the methionine adenosyltransferase (MAT2a) enzyme that directly utilizes methionine to generate the universal methyl donor S-adenosylmethionine (SAM), which is used by methyltransferases for histone and DNA modification^18^. *AHCY* encodes the adenosylhomocysteinase (AHCY) enzyme that reversibly converts the unmethylated byproduct S-adenosylhomocysteine (SAH) to generate homocysteine and adenosine^19^. *MAT2A* dependency was also found for other lineages including breast cancer, pancreatic cancer, and Ewing sarcoma (**Figure S1a**). Other cell lines demonstrating significant *AHCY* dependency were associated with immune system cancers (**Figure S1b**). In total, 71 CNS cell lines showed significant dependency for *MAT2A* (mean gene effect -1.273 ± 0.407) and 36 showed significant dependency for *AHCY* (mean gene effect - 0.680 ± 0.143), with 35 CNS cell lines exhibiting significant dependency for both genes (**Figures S2a, b**). Reactome pathway enrichment analyses were conducted using the STRING Database (https://string-db.org)^20,21^ and indicated that sulfur amino acid metabolism was significantly enriched for both *MAT2A* and *AHCY* (**Figures 1f, g**). Sulfur is derived from methionine, and its metabolism constitutes the cellular antioxidant system responsible for maintaining redox homeostasis^22^. Pathway enrichment and *MAT2A* and *AHCY* genetic dependency collectively suggest that these two genes and their encoded enzymes are candidates for targeting redox balance and cell growth in GBM.

**Figure 1.**
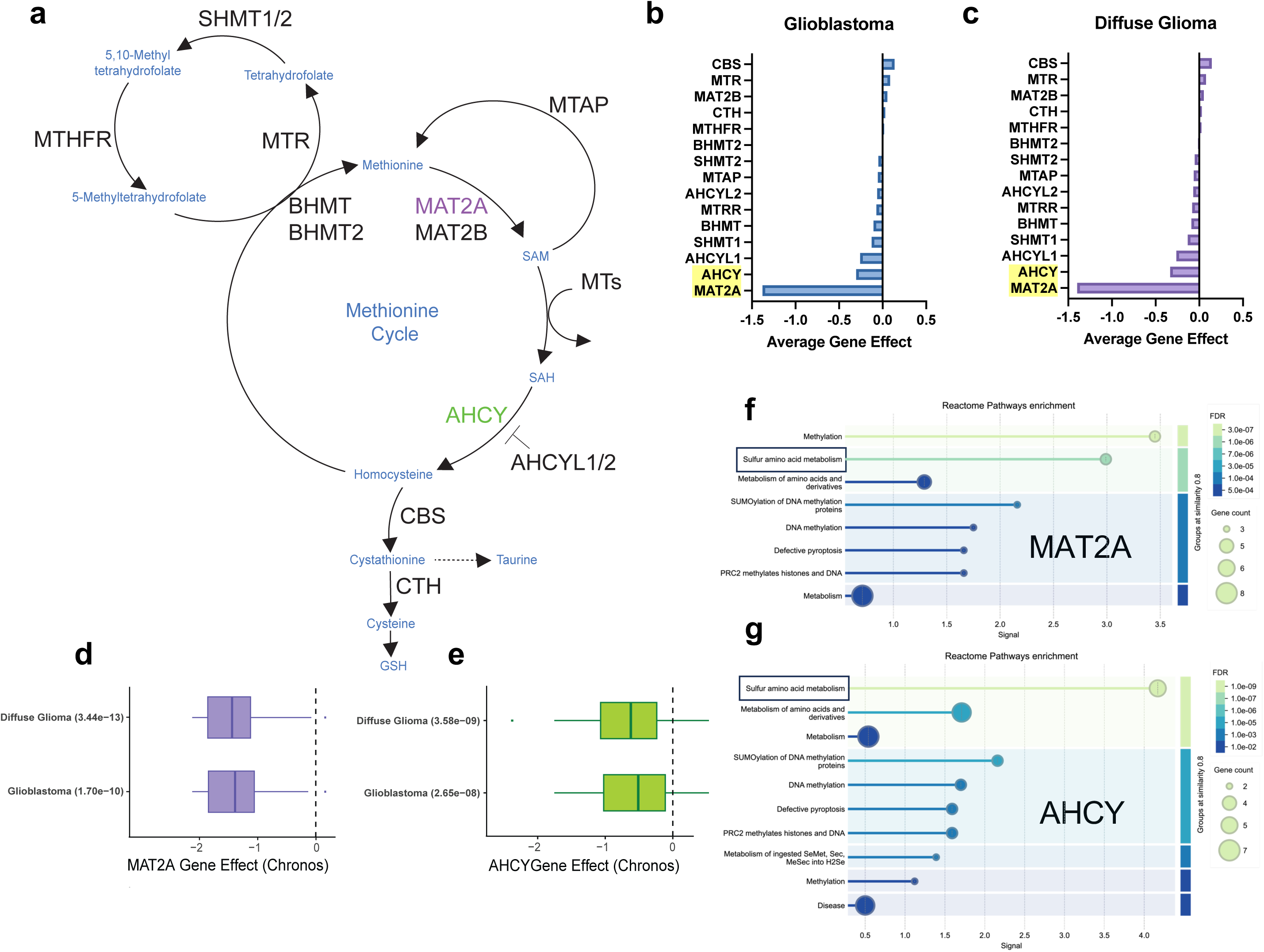
Pooled CRISPR screen data reveals methionine metabolism gene essentiality in GBM. **a** The methionine cycle contains multiple enzymes to generate methyl donor SAM, and downstream antioxidant substrates. **b, c** Average gene effect scores representative of DepMap pooled CRISPR screens for indicated genes involved in the methionine cycle in Glioblastoma and Diffuse Glioma lineages. Less than 0 indicates impaired cell growth and greater than 0 indicates enhanced cell growth. **d, e** Gene effect scores for both MAT2A and AHCY in Diffuse Glioma and Glioblastoma lineages represented as boxplots scores filtered using p-value <0.00005 according to DepMap significance. **f, g** Significantly enriched pathways from the Reactome database for both MAT2A and AHCY, reflecting high signal for sulfur amino acid metabolism, generated using the STRING database.

### siRNA-induced knockdown of *MAT2A* and *AHCY* promotes cell death and lipid peroxidation in GBM

Based on these significant genetic dependency scores and pathway enrichments, we wanted to confirm the CRISPR screen results and further evaluate the role of *MAT2A* and *AHCY* in GBM cell survival and redox balance. We used selective siRNAs targeting either *MAT2A* or *AHCY* to evaluate the effect of gene silencing on LN229 cell survival. MAT2a and AHCY protein expression was significantly reduced (siMAT2A #1 11.98 ± 4.72, p < 1×10^-4^; siMAT2A #2 44.18 ± 20.25, p= 8.8×10^-3^; siAHCY #1 30.31 ± 13.59, p= 9×10^-4^; 46.04 ± 16.53, p= 4.8×10^-3^ compared to controls) at 48 hours post-transfection (**Figure 2a; Figures S3a-f**). Flow cytometry analysis revealed that there was a significant reduction in percentage survival at 96 hours after *MAT2A* knockdown (mean 53.733 ± 19.409% dead cells) compared to control (p= 0.0199) (**Figure 2b; Figure S4c**).

**Figure 2.**
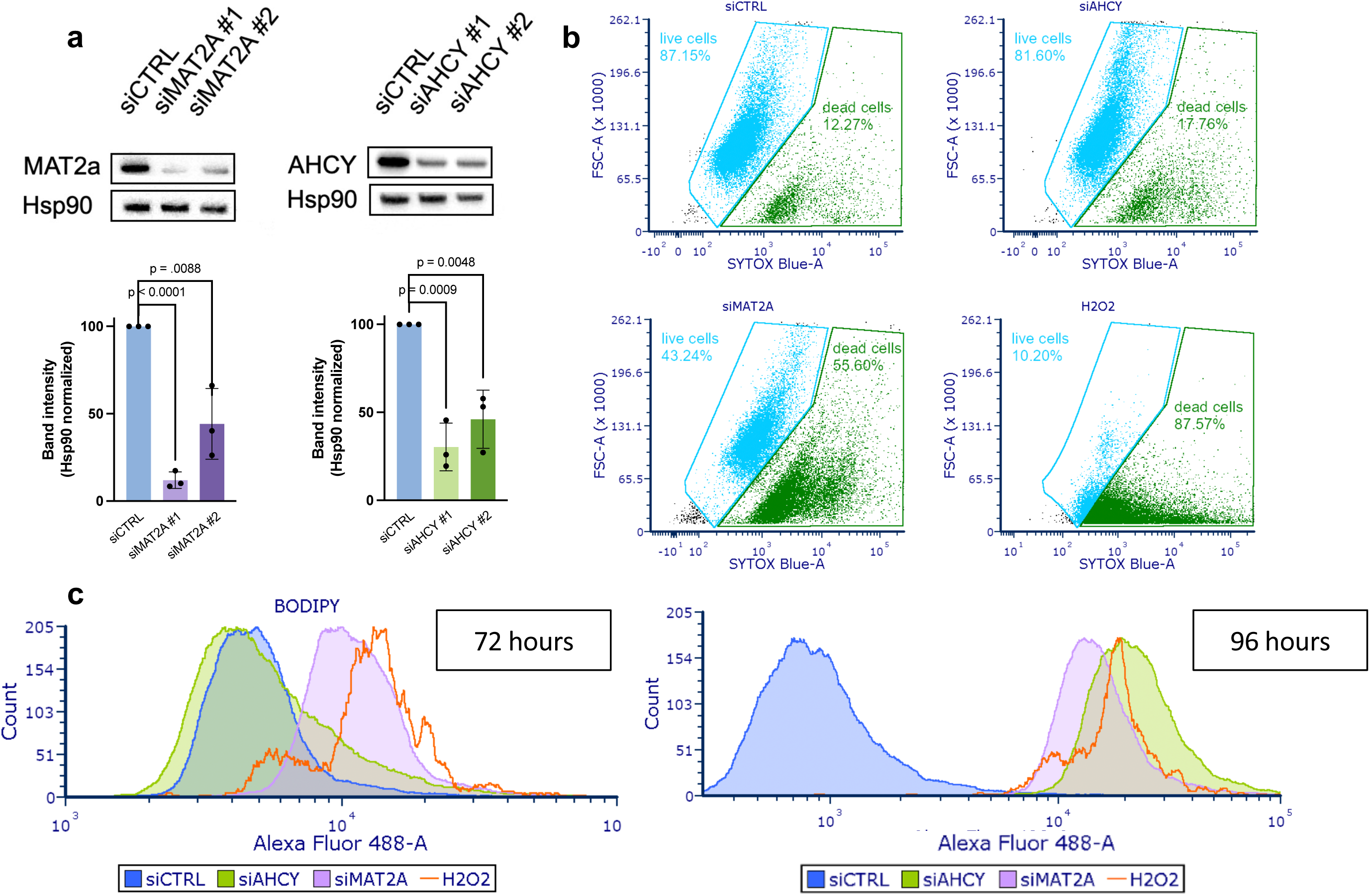
Genetic knockdown of MAT2A and AHCY induces cell death and lipid peroxidation in GBM. **a** Gene knockdown in LN229 cells 48 hours post-transfection, with heat shock protein 90 (Hsp90) as the loading control. Blots quantified using ImageJ and normalized to siCTRL band intensity. **b** Cells stained with SYTOX Blue for flow cytometry analysis of cell death 96 hours post-transfection, with hydrogen peroxide as positive control. **c** BODIPY staining for flow cytometry analysis of lipid peroxidation 72 hours and 96 hours post-transfection as indicated on the graph. Cell counts were normalized to peak intensity, with hydrogen peroxide as positive control.

Lipid peroxidation is a process that occurs at the lipid membrane due to reactive oxygen species (ROS) disrupting lipid integrity, and excess amounts lead to ferroptosis or oxidative cell death^23^. Cells rely on antioxidant protection to attenuate lipid peroxidation and prevent cell death^24^. Compared to non-malignant cells, cancer cells demonstrate higher levels of lipid peroxidation that support their growth and survival^25^. In GBM, lipid peroxidation has been observed at the invasive front of the tumor and is found to facilitate immune and therapeutic resistance^26^. However, excess lipid peroxidation leads to compromised functioning and ferroptotic cell death^27^, and this has been previously linked to methionine restriction in GBM^15^. Upon *MAT2A* knockdown, we observed an increase in lipid peroxidation (BODIPY) staining at 72 hours (**Figure 2c; Figure S4b**). After 96 hours, we observed a further increase in lipid peroxidation for *MAT2A* knockdown as well as increased lipid peroxidation staining for *AHCY* knockdown cells compared to controls (**Figure 2c; Figure S4c**). These results confirm that the GBM cell line LN229 is dependent on *MAT2A* for survival and suggest that *MAT2A* or *AHCY* are required to evade oxidative damage in GBM.

### MAT2A and AHCY knockdown compromises mitochondrial function in GBM

We sought to investigate the oxidative metabolism changes associated with *MAT2A* and *AHCY*. The primary function of mitochondria is to supply energy to cells through oxidative phosphorylation (OXPHOS)^28^. They contribute two primary sources of ROS generation during OXPHOS via the electron transport chain complexes I and III^29^. It is well appreciated that maintaining ROS at appropriate levels is necessary for normal cell function^30^, but elevated levels of ROS can be activating for cancer proliferation, invasion, and metastasis in GBM^31–33^. Therefore, we identified key parameters of mitochondrial bioenergetics following genetic knockdown. We conducted the Mitochondrial Stress Test (**Figure 3a**) using the Seahorse bioanalyzer, which can measure oxidative metabolism. Cellular respiratory parameters can be calculated based on changes in dissolved oxygen and pH in live cell media. The readouts are translated to represent oxygen consumption rate (OCR) and extracellular acidification rate (ECAR), respectively. Following the addition of electron transport chain complex inhibitors (complex V inhibitor oligomycin^34^, or complex I/III inhibitors rotenone/antimycin A^29^) or a proton uncoupler (FCCP^35^) over a set time course during the experiment, we can uncover potentially impaired components of OXPHOS in knockdown cells. Given that *MAT2A* and *AHCY* are implicated in antioxidant production^11^, we anticipated that silencing these genes would result in disrupted oxidative metabolism such that the deleterious effects of ROS accumulation would lead to reduced GBM survival.

**Figure 3.**
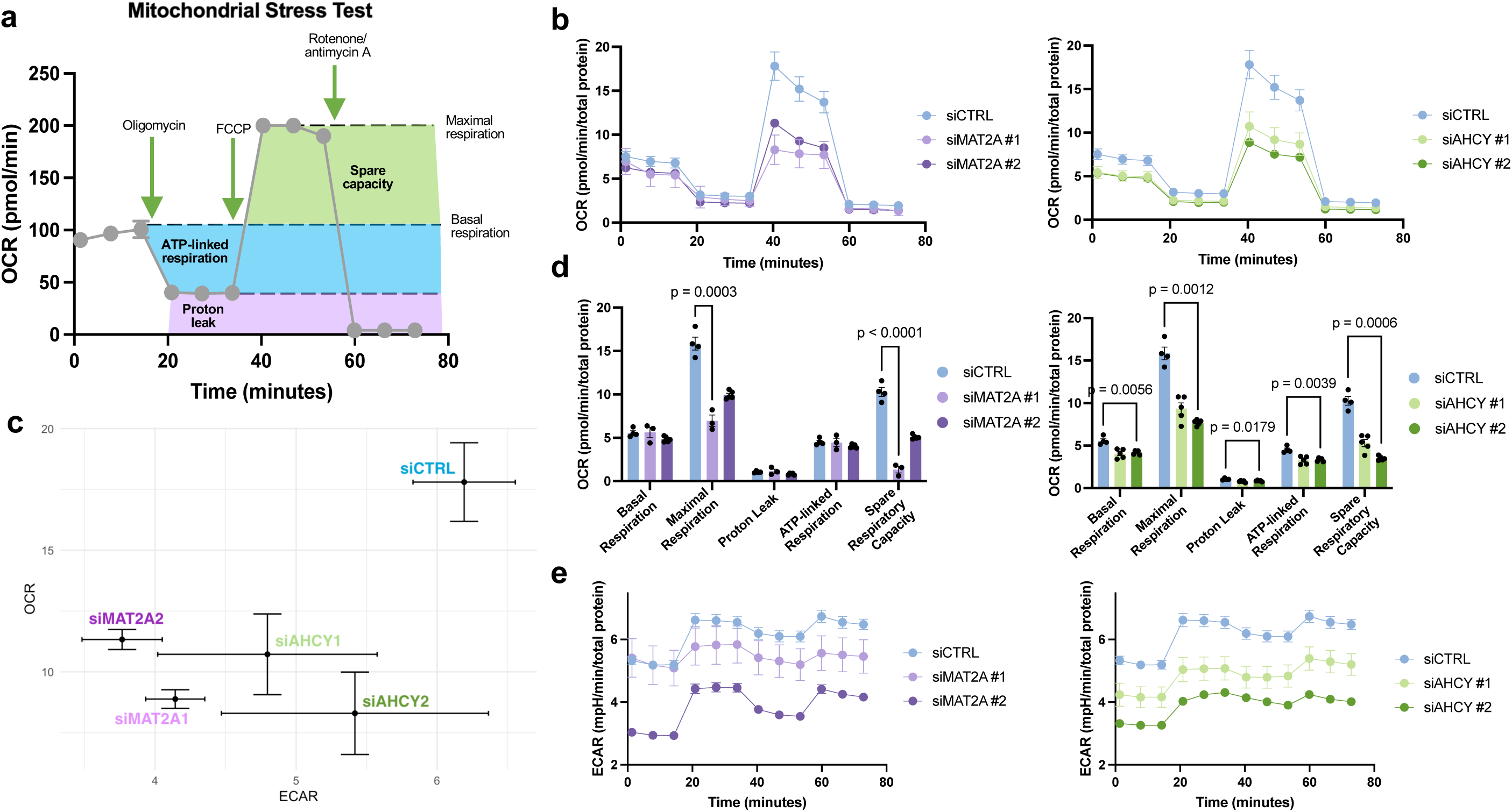
Genetic knockdown of MAT2A and AHCY induces mitochondrial dysfunction in GBM. **a** Graphic of Mitochondrial Stress Test indicating changes in oxygen consumption rate (OCR) following electron transport chain inhibitor compound injections. Highlighted regions correlate to respiratory parameters by calculated differences between OCR values. **b** Mitochondrial Stress Test OCR with LN229 cells with genetic knockdown of either MAT2A or AHCY 72 hours post-transfection. **c** Energy map with mean maximal OCR and corresponding ECAR values of genetic knockdown LN229 cells, with top right indicating energetic, bottom right indicating glycolytic, top left indicating aerobic, and bottom left indicating quiescent phenotype. **d** Quantification of respiratory parameters for MAT2A and AHCY knockdown cells compared to control. Unpaired t-test with Holm-Šídák correction was performed. **e** Mitochondrial Stress Test ECAR with LN229 cells corresponding to OCR measurements, with top right corner indicating a more energetically active state and bottom left corner indicating a more energetically inactive state.

Following FCCP injection, we found a diminished maximal oxygen consumption rate for both *MAT2A* and *AHCY* knockdown LN229 cells (**Figure 3b; Figure S5a**). This reflects a significant impairment of mitochondrial function and consequently reduced capacity to respond to energetic demands. In assessing maximal respiration rates and complimentary extracellular acidification rates, *MAT2A* knockdown showed significantly reduced rates (mean OCR 8.881 ± 0.383) compared to control (p < 1×10^-4^), as did *AHCY* knockdown (mean 10.721 ± 1.659, p= 4×10^-4^). When evaluating both OCR and ECAR simultaneously, this is depicted as a shift from a highly energetic state towards an energetically inactive state (**Figure 3c; Figure S5d**). Compared to the control condition, both si*MAT2A*-LN229 and si*AHCY*-LN229 cells exhibited significantly reduced maximal respiration (p= 3×10^-4^, 1.2×10^-3^ respectively), spare respiratory capacity (p <1×10^-4^, 6×10^-4^ respectively), and overall reduced glycolytic activity (**Figures 3d, e; Figures S5b, c**). This suggests that both *MAT2A* and *AHCY* are integral to preventing mitochondrial ROS overload to facilitate increased energy production and proliferation in GBM.

### MAT2a and AHCY inhibition impedes cell growth in GBM primary cells

To evaluate the effect of inhibiting the enzymatic activity of MAT2a and AHCY, we identified two commercially available selective inhibitors, AG-270 and Aristeromycin, to use in further experiments. AG-270 is the first-in-class oral selective MAT2a allosteric inhibitor and is currently being tested in clinical trials for lymphoma and solid tumors^36^. Aristeromycin is a naturally occurring compound first isolated from *Streptomyces citricolor* in 1968^37^. It demonstrates potent AHCY inhibition and antiviral activity^38^, and although it has not advanced into clinical development, it has been investigated preclinically in prostate cancer with considerable efficacy^39^. Patient tumors are categorized into three molecular subtypes based on their transcriptional profile: classical, mesenchymal, or proneural^40^. We treated multiple patient-derived xenograft (PDX) GBM cells designated as having a classical molecular subtype and one proneural cell type (**Table S1**) using these two compounds individually (**Figure S6a**). Two cell types, GBM6 and GBM76, were characterized as having a classical molecular subtype and were taken at initial diagnosis or upon tumor recurrence, respectively (**Table S1; Figures S6b-e**). We were interested in analyzing classical PDX cells as they demonstrated the highest MAT2A and AHCY expression across subtypes based on RNASeq, Agilent-42502A, and HG-U133A datasets from the Cancer Genome Atlas (http://gliovis.bioinfo.cnio.es/)^41^. Following 24-hour treatment with AG-270, there was a significant reduction in S-adenosylmethionine (SAM, p < 1×10^-4^ for both) as well as S-adenosylhomocysteine ( SAH, p= 4.2×10^-3^, 2×10^-4^ respectively) in both GBM6 and GBM76 compound-treated cells compared to vehicle control (**Figures 4a, b; Figures S6f, g**). Furthermore, there was a significant decrease in SAM (p < 1×10^-4^, p= 3×10^-4^ respectively) and a significant increase in SAH (p= 2.5×10^-3^, p < 1×10^-4^ respectively) following Aristeromycin treatment in either cell (**Figures 4a, b; Figures S6f, g**). These changes in metabolite levels were expected considering these inhibitors are selectively targeting the enzymes responsible for producing or consuming these specific metabolites (**Figure 4c**). To study the effect of these compounds on cell viability, cells were treated with increasing concentrations of AG-270 or Aristeromycin and evaluated based on the detection of ATP after 72 hours. Both treatments led to a dose-dependent reduction in ATP content with an EC_50_ value below 6µM for AG-270 (3.61 ± 2.14µM in GBM6; 5.56 ± 2.80µM in GBM76) and 3µM for Aristeromycin (2.05 ± 0.08µM in GBM6; 0.61 ± 0.04µM in GBM76) in both cell types (**Figures 4d, e**). Only metabolically active cells are considered viable, so reduced ATP content is indicative of reduced cell viability. These results indicate that both AG-270 and Aristeromycin effectively engage with their target enzyme and potently inhibit its function at low micromolar concentrations, leading to reduced GBM cell viability.

**Figure 4.**
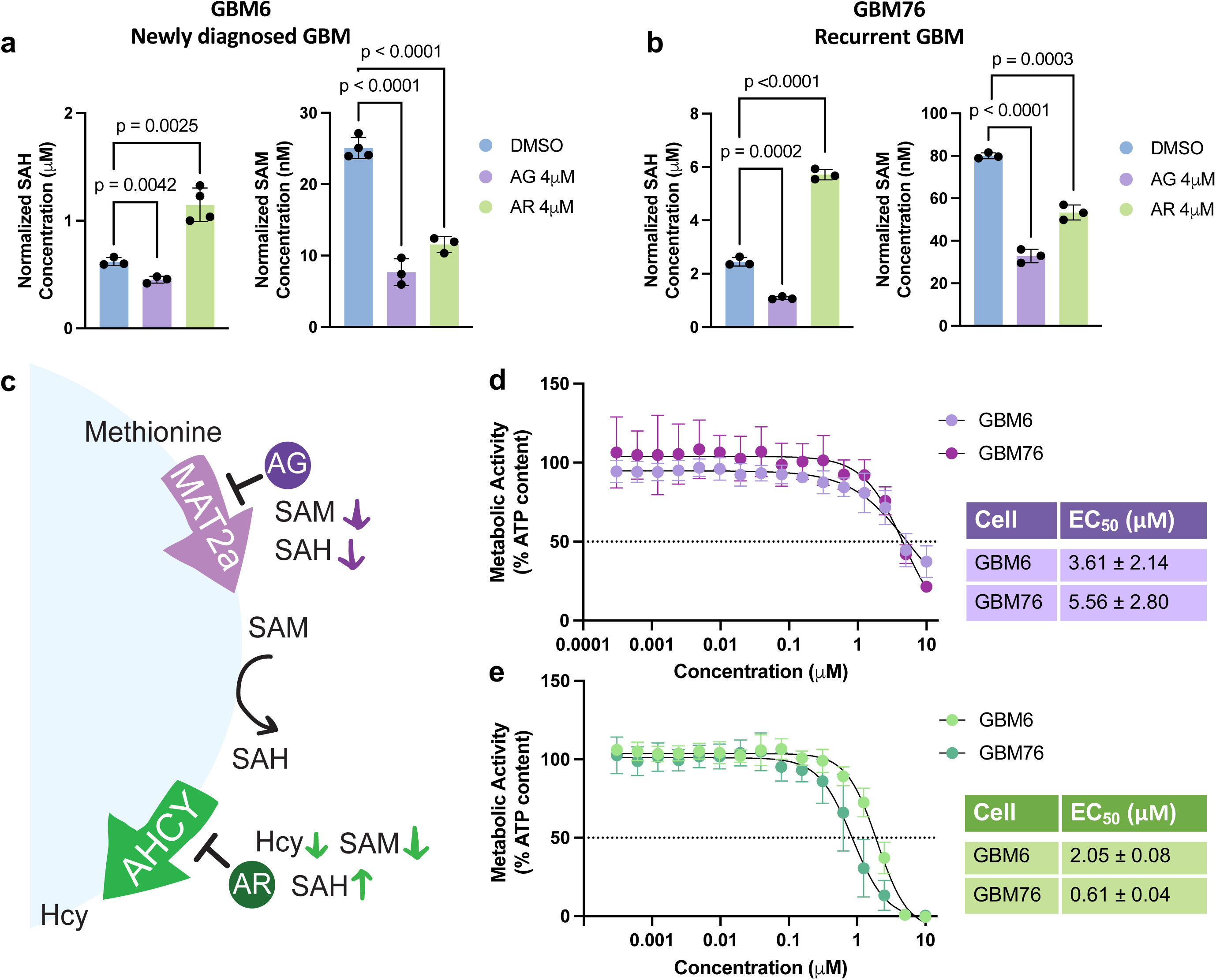
Selective MAT2A and AHCY inhibitors reduce cell viability in GBM. **a,b** SAH and SAM levels in patient-derived primary GBM cells 24 hours after 4µM of Aristeromycin (AR), 4µM of AG-270 (AG) treatment or vehicle treatment. **c** Graphic of enzyme inhibition by each respective inhibitor compound and resulting changes in metabolite levels. **d, e** Dose-response curves for AG-270 (top) and Aristeromycin (bottom) 72 hours post-treatment with corresponding EC_50_ values in both GBM6 and GBM76 cells. ATP content was measured using CellTiterGlo to indicate cell viability.

### MAT2a and AHCY inhibition compromises mitochondrial function in GBM primary cells

We sought to discover the perturbations in oxidative metabolism associated with MAT2a or AHCY inhibition. We again conducted the Mitochondrial Stress Test to evaluate mitochondrial respiration following short-term (24-hour) inhibitor treatment using the Seahorse bioanalyzer. AG-270 significantly reduced cellular respiration in GBM6 cells, with lower maximal respiration (p= 9×10^-4^) and spare respiratory capacity (p= 0.0145) (**Figures 5a, b; Figures S8a, b**). Aristeromycin significantly reduced multiple parameters of cellular respiration in GBM76 cells, including basal respiration (p= 0.0323), maximal respiration (p= 6.9×10^-3^), ATP-linked respiration (p= 0.0263), and spare respiratory capacity (p= 4.6×10^-3^) (**Figures 5d, e; Figures S8d, e**). In both cases, the inhibitors also reduced glycolytic activity according to the extracellular acidification rate relative to vehicle control (**Figures 5c, f; Figures S8c, f**). These results indicate that MAT2A or AHCY inhibition significantly reduces cellular respiration and mitochondrial function in GBM.

**Figure 5.**
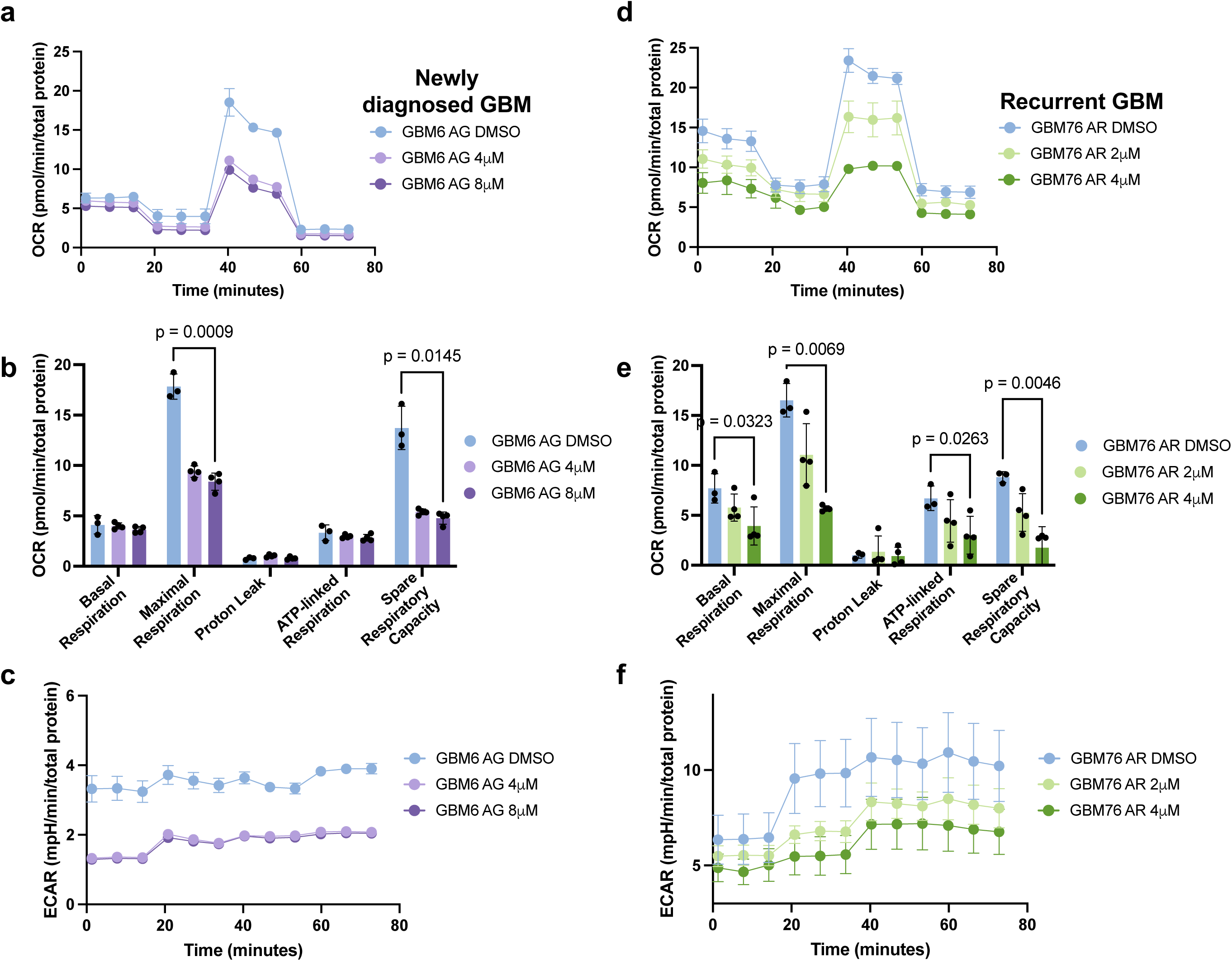
MAT2A and AHCY enzyme inhibition disrupts mitochondrial function in GBM. **a** Mitochondrial Stress Test OCR with newly diagnosed GBM6 cells treated with AG-270 72 hours prior. **b** OCR for recurrent GBM76 cells treated with Aristeromycin 72 hours prior. **c, d** Quantification of respiratory parameters corresponding to OCR measurements compared to control. Performed unpaired t-test with Holm-Šídák correction. **e, f** Mitochondrial Stress Test ECAR corresponding to OCR measurements.

### MAT2a and AHCY inhibition reduces antioxidant production and induces oxidative stress in GBM

To characterize the metabolic profile of primary GBM cells following inhibitor treatment, we conducted targeted metabolomic analysis to quantitatively measure specific groups of metabolites in these samples and potentially uncover novel associations between metabolite levels and the respective treatment conditions for GBM6 and GBM76 cells^42^. We wanted to assess how inhibition of either MAT2a or AHCY impairs oxidative metabolism. We used a panel of 366 biochemically annotated one-carbon metabolites and TCA cycle metabolites as our internal standard to identify the metabolites within our samples. This panel allowed us to measure not only metabolites that are directly involved in the methionine cycle but also those that are broadly involved in one-carbon metabolism-nucleotide biosynthesis, lipid metabolism and transport, polyamine synthesis, B vitamin metabolism, neuroprotective programs^43^- and the TCA cycle-highlighting key factors for mitochondrial function^44^. To ensure that serum supplementation would not artificially influence cellular metabolism and thus skew intracellular metabolite levels, we cultured GBM6 and GBM76 cells using serum-free media for our metabolomics analysis. Groups were evaluated using principal component analysis (PCA) and were found to cluster by treatment condition (**Figures 6a, b; Figure S10a**). Both GBM6 and GBM76 cells treated with Aristeromycin demonstrated a significant increase in S-adenosylhomocysteine (log2FC 0.98, p= 7.5×10^-3^ for GBM6; log2FC 2.17, p= 1.4×10^-3^ for GBM76, **Figure 6d; Figure S10b**), as expected from previous analysis (**Figures 4a, b; Figures S6f, g**). Several other metabolites were significantly reduced or increased commonly across different treatment groups and cell types (**Figures S11a-c; Table S2**).

**Figure 6.**
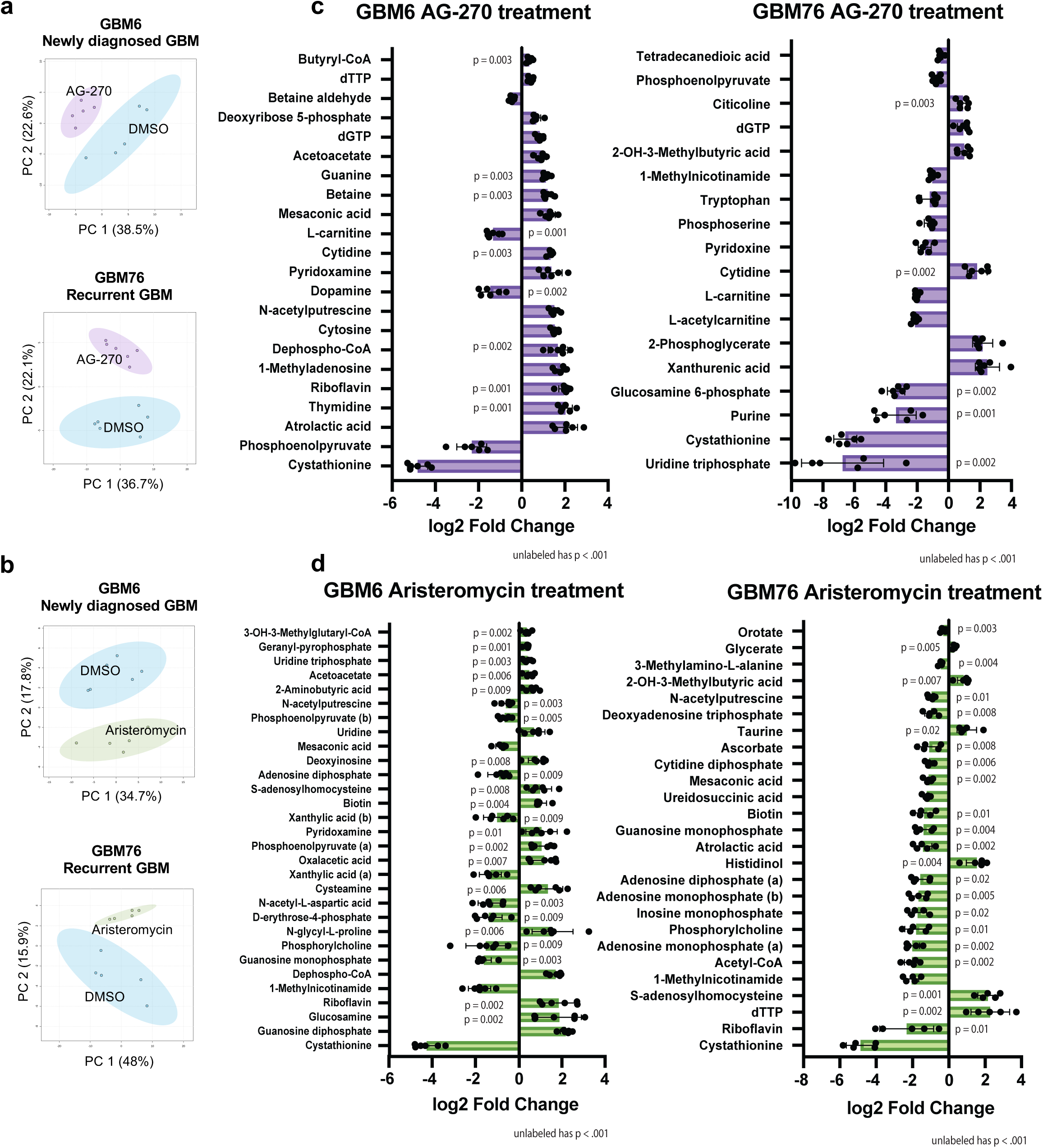
GBM exhibits oxidative stress and compromised lipid metabolism upon MAT2A and AHCY inhibition. **a,b** Principal component analysis of GBM6 and GBM76 cells grown in serum free media comparing DMSO to AG-270 (a) or Aristeromycin (b) treatment. Analyzed after 24 hours using LC-MS. Sample peak intensities were normalized using log transformation and Pareto scaling. **c,d** Significantly reduced and increased metabolites in GBM6 and GBM76 cells treated with AG-270 (c) and Aristeromycin (d) compared to control using unpaired t-test with a p-value of .05. **(b)** indicates the alkaline metabolite while **(a)** indicates the acidic metabolite upon ionization.

One metabolite in particular that is downstream of methionine metabolism is cystathionine, a sulfur-containing molecule, which is implicated in redox homeostasis as a substrate for glutathione synthesis^45^. Cystathionine was markedly depleted in all treatment groups (log2FC -4.82, p= 2.02×10^-6^ for GBM6 AG-270; log2FC -4.28, p= 3.01×10^-7^ for GBM6 Aristeromycin; log2FC -6.56, p= 9.13×10^-6^ for GBM76 AG-270; log2FC -4.85, p= 6×10^-4^, **Figures 6c, d; Figures S11a, b**). The remaining top significant metabolites that were reduced following AG-270 treatment in both cell types were L-carnitine (log2FC -1.33, p= 1.2×10^-3^ for GBM6; log2FC -2.02, p= 6.27×10^-5^ for GBM76), which transports long chain fatty acids into the mitochondria to be oxidized for energy^46^ and is associated with ROS scavenging and enhancing antioxidant capacity^47^, and phosphoenolpyruvate (log2FC -2.31, p= 4×10^-4^ for GBM6; log2FC - 0.76, p= 6×10^-4^ for GBM76), a high-energy metabolic intermediate that supports cancer proliferation and balances ROS levels^48,49^ (**Figure 6c, Figure S11a**). Among the metabolites that were increased in both cell types following AG-270 treatment were cytidine (log2FC 1.33, p= 3×10^-3^ for GBM6; log2FC 1.84, p= 1.6×10^-3^ for GBM76), a nucleoside and an important substrate for mitochondrial biogenesis^50^, and cholesteryl sulfate (log2FC 1.96, p= 0.0158 for GBM6; log2FC 1.12, p= 0.020 for GBM76), which inhibits cholesterol synthesis^51^ and upregulates antioxidant capacity in normal astrocytes^52^, and 1-methyladenosine (log2FC 1.79, p= 6.01×10^-5^ for GBM6; log2FC 1.12, p= 3.6×10^-3^ in GBM76), which behaves as an mRNA modification that is upregulated during neuronal oxidative stress damage^53^ (**Figure 6c, Figure S11a**). The remaining metabolites that were significantly reduced upon Aristeromycin treatment in both cell types included phosphorylcholine (log2FC -1.55, p= 8.9×10^-3^ for GBM6; log2FC - 1.80, p= 0.0144 for GBM76), which is involved in lipid homeostasis and associated with GBM proliferation^54^, 1-methylnicotinamide (MNA, log2FC -1.83, p= 7.75×10^-6^ for GBM6; log2FC -2.04, p= 4.17×10^-6^ for GMB76), a downstream product of methionine metabolism that coordinates energy metabolism^55^, guanosine monophosphate (log2FC -1.53, p= 0.0225 for GBM6; log2FC - 1.39, p= 4.1×10^-3^ for GMB76), a precursor for vasodilation and neurotransmission signaling and is associated with recurrence in GBM^56^, mesaconic acid (log2FC -0.84, p= 9.54×10^-5^ for GBM6; log2FC -1.12, p= 2.1×10^-3^ for GMB76), a fatty acid associated with reduced inflammatory response in the brain^57^, and N-acetylputrescine (log2FC -0.62, p= 2.9×10^-3^ for GBM6; log2FC - 0.95, p= 0.0138 for GMB76), which is involved in fatty acid oxidation and is higher in GBM compared to control patients^58^ (**Figure 6d, Figure S11b**). Other than SAH, oxalacetic acid (log2FC 1.17, p= 6.7×10^-3^ for GBM6; log2FC 0.45, p= 0.0320 for GBM76), shown to promote brain mitochondrial biogenesis and reduce neuroinflammation^59^; and glycerate (log2FC 0.23, p= 0.0305 for GBM6; log2FC .32, p= 4.5×10^-3^ for GBM76), a product of glycerol oxidation that acts as a metabotoxin at sufficiently high levels^60^, were the two significantly upregulated metabolites for both cell types treated with Aristeromycin (**Figure S11a**). 3-Hydroxy-3-methylglutaryl-CoA, the metabolite whose conversion instigates cholesterol synthesis^61^, was selectively increased in GBM6 cells treated with Aristeromycin (log2FC 0.36, p= 0.0016, **Figure 6d**). Collectively, these results suggest that MAT2a or AHCY inhibition compromises GBM cellular metabolism by disrupting antioxidant production, fatty acid transport, cell membrane integrity, nucleotide synthesis, and other related pro-tumorigenic programs.

## Discussion

In this study, we demonstrate that siRNA-mediated depletion and pharmacologic inhibition of MAT2a or AHCY hindered GBM cell survival. Cell viability was reduced in *MAT2A* knockdown cells, and both knockdown conditions exhibited increased lipid peroxidation. Knockdown of *MAT2A* or *AHCY* led to a significant reduction in cellular respiration, as evidenced by a decrease in oxygen consumption and extracellular acidification rates and a shift towards an energetically inactive state. Pharmacological inhibition of both MAT2a and AHCY resulted in reduced cell viability at low concentrations. Cellular respiration was reduced in both GBM6 and GBM76 cells following MAT2a and AHCY inhibition, respectively. Inhibitor-treated cells also exhibited increased pro-oxidative markers, reduced antioxidative markers, and compromised energy production due to oxidative damage. These results suggest that restricting methionine metabolism by these two targets reduces antioxidant capacity, that enzyme inhibition potency may be predicated on proliferative phenotype, and that the shift in metabolic profile highlights a vulnerability thwarting invasion, energy production, and treatment resistance in GBM.

Notably, the DepMap data revealed that LN229 cells showed significant dependency only for *MAT2A* and not *AHCY* (**Figure S2b**). Thus, our observation that *AHCY* knockdown does not significantly affect LN229 cell survival is expected. Even still, there was evidence of increased lipid peroxidation (**Figure 2c; Figure S4c**) and a significant reduction in spare respiratory capacity in these cells following *MAT2A* or *AHCY* knockdown (**Figures 3a, b; Figures S5 a, b**). Mitochondrial spare respiratory capacity, or uncoupling control ratio, refers to the cell’s ability to adapt to cellular stress and manage increased respiration when met with greater energy demands^62^. Spare respiratory capacity is therefore highly dependent on redox regulation^63^ and is significantly reduced by oxidative stress through several different mechanisms involving ROS accumulation and reduced antioxidant stores^64–66^. Enzyme inhibitor treatment also significantly reduced spare respiratory capacity in both GBM6 and GBM76 cells (**Figures 5a, b, d, e; Figures S8a, b, d, e**) indicating they were less amenable to energetic demands and more vulnerable to oxidative stress. Impaired redox regulation not only reduced spare capacity but also impeded glycolytic respiration (**Figures 5c, f; Figures S8c, f**) thereby hindering cellular respiration and energy production altogether. Therefore, MAT2a and AHCY are both implicated in antioxidant capacity and protection against oxidative stress in GBM.

There was an apparent correlation between enzyme inhibition and reduced cell viability for the two cell types; GBM76 cells appear to be more sensitive to AHCY inhibition while GBM6 cells appear to be more sensitive to MAT2a inhibition. GBM76 cells exhibited a more significant increase in SAH levels and had a lower EC_50_ following Aristeromycin treatment (**Figures 4a, b, d**). GBM76 cells treated with Aristeromycin also demonstrated a greater fold change increase in SAH based on metabolomics analysis (**Figure 6d**). By contrast, the MAT2a inhibitor AG-270 led to a more marked reduction in SAM levels in GBM6 cells (**Figures 4a, b**) and had a lower EC_50_ in GBM6 cells compared to GBM76 cells (**Figure 4c**). Furthermore, there was a corresponding reduction in spare respiratory capacity and glycolytic activity unique to GBM76 cells treated with Aristeromycin (**Figures 5d-f; Figures S8d-i**) as well as for GBM6 cells treated with AG-270 (**Figures 5a-c; Figures S8a-c, j-l**). GBM6 and GBM76 cells displayed distinctly abundant metabolites in their baseline metabolomic profiles (**Figures S9a-c**). When we compared the basal metabolic rates of GBM6 and GBM76 cells, it was evident that the GBM76 cells exhibited significantly higher basal respiration and reduced spare capacity (**Figures S7a, b**). Reduced spare capacity of cancer cells at baseline may indicate that the cells are highly proliferative^67^, therefore GBM76 cells appeared to be siphoning a greater portion of their spare capacity to support aggressive proliferation. However, upon AHCY inhibition, GBM76 cells exhibited reduced spare capacity, basal respiration (**Figures 5a, b, d, e; Figures S8a, b, d, e**), and viability (**Figure 5e**), suggesting that reduced spare capacity in response to treatment is not reflective of increased proliferation. This points to the relevance of an aggressive proliferative phenotype in the potency of either MAT2a or AHCY inhibition in GBM.

A recent study identified the downstream methionine substrate cystathionine to be the most highly abundant metabolite in the invasive tumor edge fraction compared to the tumor core fraction, implicating antioxidant metabolism in GBM invasion^26^. In our study, cystathionine was most depleted consistently across all treatment groups (**Figures 6c, d**). This suggests that both GBM cell viability and invasion are associated with MAT2a or AHCY inhibition. Further studies need to be performed to determine how MAT2a or AHCY inhibition impacts GBM cell invasion. GBM76 cells treated with Aristeromycin had significantly reduced levels of alkaline biotin and riboflavin (**Figure S11b**). Biotin and riboflavin are important B vitamins that facilitate carboxylation and redox reactions in the brain, respectively^68^. Riboflavin is critical for mitochondrial aerobic respiration while biotin is necessary for mitochondrial fatty acid oxidation and gluconeogenesis^69^. A previous study highlighted that biotinylation is an important modification in glioma stem cells (GSCs) and disrupted biotin distribution leads to cholesterol depletion, impaired OXPHOS, disrupted GBM proliferation, and reduced invasiveness^70^. This indicates that this post-transcriptional modification may be diminished in GMB76 cells and upregulated in response to GBM6 methionine pathway inhibition. L-carnitine was more abundant in GMB6 cells at baseline (**Figure S9c**) and was significantly depleted in both cell types treated with AG-270 (**Figure 11b**). GBM relies on L-carnitine for antioxidant enzyme activity^71^; it is more highly abundant in both newly diagnosed and recurrent GBM compared to normal brain tissue^72^; and supplementation mitigates cell death from TMZ or hydrogen peroxide^72^. This suggests that MAT2a inhibition may promote GBM re-sensitization to standard-of-care treatment through diminished antioxidant capacity.

While our findings highlight the importance of MAT2a and AHCY in antioxidant metabolism, other potential tumorigenic mechanisms may be involved that were not explored. Pancreatic ductal adenocarcinoma is characterized by severe hypoxia, and recent evidence suggests that the hypoxic environment specifically protects MAT2a from degradation to facilitate enhanced nutrient cycling^73^. In breast cancer, the cytoplasmic:nuclear MAT2a protein expression ratio is correlated with invasiveness and poor overall survival, and this is expected to be relevant to a methylthioadenosine phosphorylase(MTAP)-deficient phenotype^74^. Previous work probing MAT2a in H3K27 mutant glioma has demonstrated that MAT2a expression tends to be higher in GBM and other high-grade gliomas compared to diffuse intrinsic pontine glioma (DIPG), and yet, even low levels of MAT2a expression are critical for DIPG survival independent of MTAP depletion^75^. This suggests that MAT2a expression is critical for high-grade glioma survival, is involved in hypoxia adaptation, and plays an important role in cell invasiveness and tumor aggressiveness. Inhibition of epigenetic regulator EZH2 reduced MAT2a and AHCY expression and promoted DNA damage and apoptosis in multiple myeloma cells^76^, indicating that methionine metabolism may be epigenetically regulated to help sustain tumor survival. Whole exome sequencing demonstrated significant upregulation and amplification of *AHCY* in drug-resistant B cell lymphoma that persists even after removal of AHCY inhibitor DZNeP^77^. This indicates that AHCY activation is a key mechanism of metabolic reprogramming to evade treatment insult and sustain tumor progression. Homocysteine, a product of AHCY enzymatic activity^19^, appears to be much higher in children with acute lymphoblastic leukemia compared to controls^78^. This evidence highlights the relevance of AHCY in immunosuppression and immune signaling in cancer.

The inhibitor compounds we used in our analysis show great promise for neuro-oncological development. Upon discovery of AG-270, researchers determined that there were no concerning off-target effects based on supporting pharmacological testing in vitro and it was well tolerated at SAM-reducing exposure in vivo^36^ although it lacks sufficient proof of ability to penetrate the blood-brain barrier. Aristeromycin analogs have been synthesized in recent years, one of which has a considerable therapeutic window as demonstrated by the calculated effective concentration to inhibit antiviral activity as well as the cytotoxic concentration to inhibit normal cell replication^79^. There is a need for future studies to establish a therapeutic window for the use of these inhibitors in the brain.

In conclusion, targeting methionine metabolism via MAT2a or AHCY inhibition is a possible avenue to arrest cancer progression and improve outcomes for GBM patients. We found that targeting these enzymes leads to compromised antioxidant capacity, reduced mitochondrial function, and cell death, as evidenced by reduced cellular respiration and reduced levels of antioxidant markers. Future studies are needed to better elucidate this mechanism and to develop clinical candidates for these inhibitors with appropriate efficacy and safety profiles.

## Methods

### Cell lines

Human glioblastoma cell line LN229 was purchased from ATCC. Patient-derived xenograft cells were acquired from Dr. Jann Sarkaria (Mayo Clinic, Rochester, MN). Briefly, cells were cultured in cell culture-treated flasks (CellTREAT) in Dulbecco’s modified Eagle’s medium (DMEM) and Ham’s F-12 nutrient mixture (F12) (Gibco; Thermo Fisher Scientific, Inc., cat. no. 11320033) supplemented with 10% fetal bovine serum (FBS) (Gibco, Thermo Fisher Scientific, Inc., cat. no. A5256801) and 100U penicillin and 100 mg/ml streptomycin (Gibco, Thermo Fisher Scientific, Inc., cat. no. 10099141). Serum-free stem cells were maintained in DMEM:F12 supplemented with neuronal cell supplement (StemCell Technologies, cat. no. 05711), 200mM L-glutamine (Corning, cat. no. 25005CI), human FGF and human EGF supplement (Thermo Fisher Scientific, Inc., cat. nos. PHG0261 & PHG0311), as well as penicillin/streptomycin (Thermo Fisher Scientific, Inc.). All cultures were maintained in a humidified incubator at 37°C with 5% CO_2_.

### Antibodies and siRNA constructs

MAT2a antibody (cat. no. PA5-115550) and AHCY antibody (cat. no. MA5-42797) were purchased from Thermo Fisher. Two Silencer^TM^ Validated siRNAs along with an ON-TARGET Plus Human siRNA smartpool (Horizon Discovery) for each gene of interest were purchased (Invitrogen, Thermo Fisher).

### DepMap Analysis

Methionine pathway genes perturbation effects were captured by generated CRISPR Chronos gene dependency scores associated with a significant p-value for glioblastoma, diffuse glioma and CNS tumor lineages. These scores and p-values were collected and sorted for each cell lineage, and effect scores were averaged for each gene target.

### siRNA transfection

LN229 cells were seeded in 6-well plates at a density of 2.4×10^5^ cells per well (or half for a 96-hour post-transfection incubation experiment). After 24 hours, once they had reached 70% confluency, cells were transfected with 100nM of each siRNA using Lipofectamine RNAiMax reagent (Invitrogen; Thermo Fisher Scientific, Inc., cat. no. 13778150). After 48 hours, cells were collected for RNA extraction, protein extraction, or flow analysis.

### Western blot

Cell lysates were generated by mechanical disruption using lysis buffer (Cell Signaling Technology, cat. no. 9803) with a protease inhibitor cocktail (Roche, Millipore Sigma, cat. no. 4693124001) and phosphatase inhibitor cocktail (Sigma Aldrich, cat. no. P5726) in deionized water. Electrophoresis was performed using a precast SDS-PAGE gel, electrophoresis chamber and power supply for 120 minutes, and subsequent semi-dry transfer with the Trans-Blot Turbo Transfer System and reagents (BioRad Laboratories, Inc.). Membrane was stained with ponceau, washed with TBST, then blocked for five minutes with EveryBlot blocking buffer (BioRad, cat. no. 12010020). Blot was then incubated in primary antibody solution for 1.5 hours, washed three times with TBST, incubated in secondary antibody solution for 1.5 hours, then washed another three times and imaged using the ChemiDoc MP Imaging System (BioRad Laboratories, Inc.).

### Flow cytometry

LN229 cells were transfected at 0 hours and the media was replaced after 72 hours. 96 hours post-transfection, media was added back to the respective wells and BODIPY (Thermo Fisher Scientific, Inc., cat. no. D3922) was added directly to the well at a concentration of 2µM. Cells were incubated for 15 minutes at 37°C and protected from light. Cells were collected and spun down along with floating cells in the media, and were washed sufficiently with PBS. Cells were finally stained with SYTOX Blue and prepared for flow analysis by the Georgetown University Core Facility. Intact cells were gated for final analysis and normalized to peak intensity and number of events per sample.

### Mitochondrial stress test

Cells were plated in a Seahorse microplate at a density of 1.8×10^4^ cells per well for 4-8 hours. Cells were subsequently treated with different concentrations of AG-270 and Aristeromycin, or DMSO vehicle treatment. After 24 hours, the cell media was exchanged with Seahorse assay media, and compound was added to the ports: 1.5µM oligomycin in port A, 1µM trifluoromethoxy carbonylcyanide phenylhydrazone (FCCP) in port B, and 0.5µM rotenone and antimycin A in port C, with port D empty. Cells were kept in a non-CO2 incubator at 37°C for an hour before beginning the assay. Using the Seahorse XFe96 bioanalyzer (Agilent Technologies), oxygen consumption rate and extracellular acidification rate were measured on an interval after each injection to obtain twelve readings for each well over the course of the experiment. The injection compounds were used from the Mitochondrial Stress Test Kit (Agilent) along with the corresponding protocol. In brief, cells were measured at baseline, then 1µM oligomycin was injected into each well to inhibit ATP synthase, so the difference from baseline reflects oxygen consumption due to ATP respiration. Then 1.5µM FCCP was injected into each well, which is a proton uncoupler to allow maximal respiration, and therefore the difference from baseline reflects the spare respiratory capacity of the cells. Finally, 0.5µM rotenone and antimycin A were added to inhibit Complex I and III, which extinguishes cellular respiration completely to reflect proton leakage in the electron transport chain. Each experiment was normalized to total protein in each well using the Pierce BCA Protein Assay Kit (Thermo Fisher Scientific, Inc.).

### Metabolic activity viability assay

CellTiter-Glo® Luminescent Cell Viability Assay kit (Promega) was used to determine the number of viable cells following inhibitor compound treatment. Cells were plated in white bottom 96-well plates at a density of 3×10^4^ cells per well in 100µL complete media. Each plate was treated with a serial dilution of 16 concentrations, along with a negative control (.2% DMSO or less) and a positive control (Bortezomib). After 72 hours, plates were removed from incubation and brought to ambient temperature for 30 minutes. 100µL of reagent was then added to each well, the plate was shaken for 30 seconds to mix thoroughly, then incubated for 10 minutes before acquiring measurements using the CLARIOstar Plus Microplate Reader.

### Targeted Metabolomics

Targeted analysis was performed by the Georgetown University Core Facility. Samples were prepared and run using the QTRAP® 5500 LC-MS/MS System (Sciex) to quantitate 332 endogenous molecules. Results were normalized to internal standards and processed using MultiQuant 3.0.3 (Sciex). Detailed information regarding sample preparation and experimental execution can be found in Supplementary Data 3.

### Metabolomics Enrichment Analysis

Pairwise comparisons between vehicle or small molecule treatment for both cell types was performed using Metaboanalyst Software. Using single-factor statistical analysis, we normalized by logarithmic data transformation and Pareto scaling. All figures were subsequently generated including the PCA plot, volcano plot, and heatmap.

### Immunoassays for metabolite detection

The experiments were conducted in accordance with each respective manual provided using approximately 1×10^6^ cells per replicate. The SAH (cat. no. MET-5151) and SAM (cat. no. MET-5152) ELISA Kits were both purchased from Cell Biolabs, Inc. (San Diego, CA). Each experiment was normalized to total protein in each sample using the Pierce BCA Protein Assay Kit (Thermo Fisher Scientific, Inc.).

### Statistical analysis

Dose-response curve data is represented in the form of mean ± standard of means (SEM). All other data is represented in the form of mean ± standard deviation (SD). GraphPad Prism Software version 10.2.1 was utilized to perform all regression analysis, paired t-tests, and one-way analysis of variance (ANOVA). All experiments were conducted three times using three or more technical replicates or in agreement with assay instructions for statistical power.

This article contains supporting information.

## Supporting information

Targeted metabolomics panel

Metabolomics methods

Metabolomics data

Fold change metabolite calculations

Comparisons between metabolite groups

Compound data

GBM76 R analysis

GBM6 R analysis

All Seahorse Data

Statistics

## Acknowledgments

We would like to thank members of the Ayad laboratory for helpful discussions. We would like to acknowledge the FACS core and the Metabolomics Shared Resource Core at Georgetown University. We would like to thank Dr. Amrita Cheema and Dr. Rebecca Riggins for helpful discussions. Research reported in this publication was supported by BellRinger at Lombardi Comprehensive Cancer Center and the National Center for Advancing Translational Sciences of the National Institutes of Health under Award Number TL1TR001431 to ECR. The content is solely the responsibility of the authors and does not necessarily represent the official views of the National Institutes of Health.

The authors declare that they have no conflicts of interest with the contents of this article.

## Abbreviations

GBM: glioblastoma
MAT2a: methionine adenosyltransferase 2A
MAT2A: gene encoding methionine adenosyltransferase 2A
AHCY: adenosylhomocysteinase/ gene encoding adenosylhomocysteinase
MTAP: methylthioadenosine phosphorylase
DIPG: diffuse intrinsic pontine glioma
DMEM: Dulbecco’s modified Eagle’s medium
ANOVA: one-way analysis of variance

