## Supplemental Tables for "MAT2a and AHCY inhibition disrupts antioxidant metabolism and reduces glioblastoma cell survival"

### Slide 1
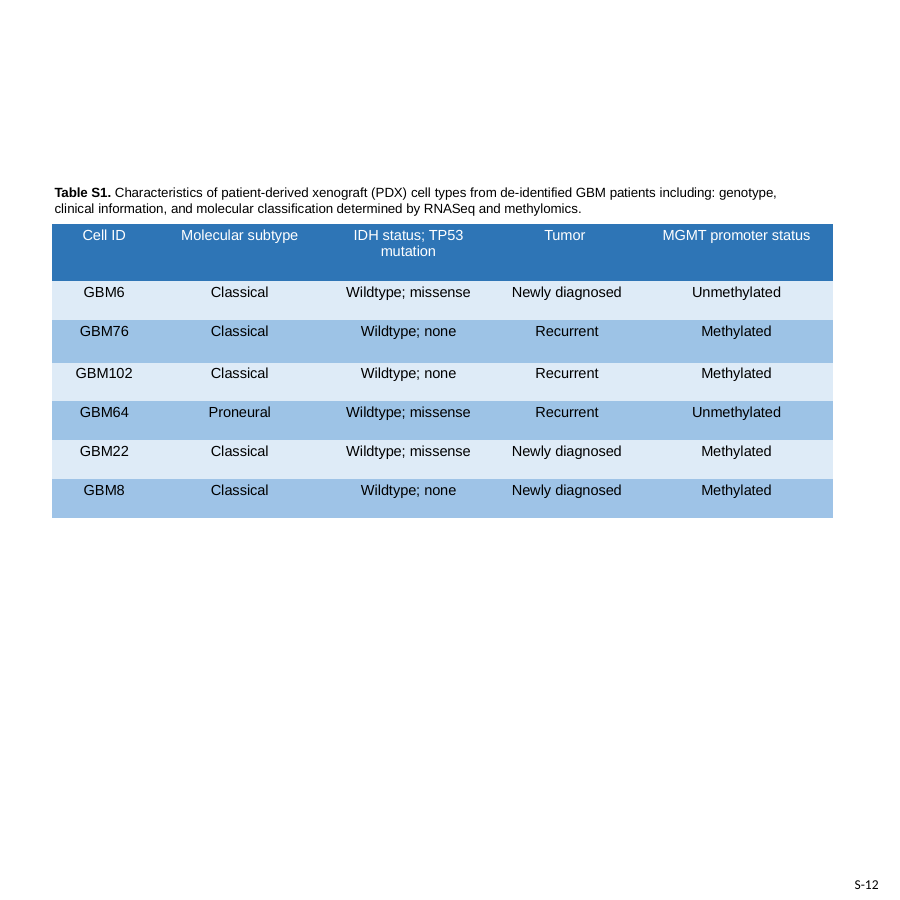

Table S1. Characteristics of patient-derived xenograft (PDX) cell types from de-identified GBM patients including: genotype, clinical information, and molecular classification determined by RNASeq and methylomics.
| Cell ID | Molecular subtype | IDH status; TP53 mutation | Tumor | MGMT promoter status |
| --- | --- | --- | --- | --- |
| GBM6 | Classical | Wildtype; missense | Newly diagnosed | Unmethylated |
| GBM76 | Classical | Wildtype; none | Recurrent | Methylated |
| GBM102 | Classical | Wildtype; none | Recurrent | Methylated |
| GBM64 | Proneural | Wildtype; missense | Recurrent | Unmethylated |
| GBM22 | Classical | Wildtype; missense | Newly diagnosed | Methylated |
| GBM8 | Classical | Wildtype; none | Newly diagnosed | Methylated |
S-12

### Slide 2
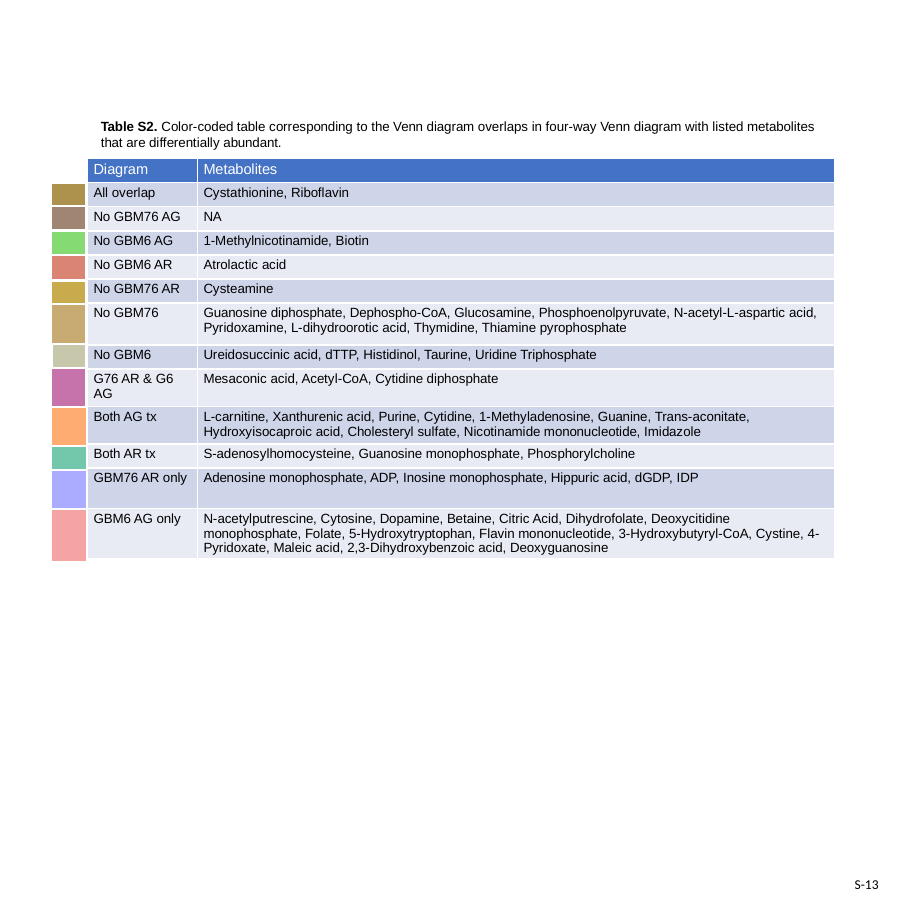

Table S2. Color-coded table corresponding to the Venn diagram overlaps in four-way Venn diagram with listed metabolites that are differentially abundant.
| Diagram | Metabolites |
| --- | --- |
| All overlap | Cystathionine, Riboflavin |
| No GBM76 AG | NA |
| No GBM6 AG | 1-Methylnicotinamide, Biotin |
| No GBM6 AR | Atrolactic acid |
| No GBM76 AR | Cysteamine |
| No GBM76 | Guanosine diphosphate, Dephospho-CoA, Glucosamine, Phosphoenolpyruvate, N-acetyl-L-aspartic acid, Pyridoxamine, L-dihydroorotic acid, Thymidine, Thiamine pyrophosphate |
| No GBM6 | Ureidosuccinic acid, dTTP, Histidinol, Taurine, Uridine Triphosphate |
| G76 AR & G6 AG | Mesaconic acid, Acetyl-CoA, Cytidine diphosphate |
| Both AG tx | L-carnitine, Xanthurenic acid, Purine, Cytidine, 1-Methyladenosine, Guanine, Trans-aconitate, Hydroxyisocaproic acid, Cholesteryl sulfate, Nicotinamide mononucleotide, Imidazole |
| Both AR tx | S-adenosylhomocysteine, Guanosine monophosphate, Phosphorylcholine |
| GBM76 AR only | Adenosine monophosphate, ADP, Inosine monophosphate, Hippuric acid, dGDP, IDP |
| GBM6 AG only | N-acetylputrescine, Cytosine, Dopamine, Betaine, Citric Acid, Dihydrofolate, Deoxycitidine monophosphate, Folate, 5-Hydroxytryptophan, Flavin mononucleotide, 3-Hydroxybutyryl-CoA, Cystine, 4-Pyridoxate, Maleic acid, 2,3-Dihydroxybenzoic acid, Deoxyguanosine |
S-13
