## Supplementary material for "MAT2a and AHCY inhibition disrupts antioxidant metabolism and reduces glioblastoma cell survival": Metabolomics methods

**Materials and methods**

All LC-MS grade solvents including acetonitrile and water were purchased from Fisher Optima grade, Fisher Scientific. High purity formic acid (99%) was purchased from Thermo-Scientific. Debrisoquine and 4-nitrobenzoic acid and 2-hyroxyglurate (2-HG) were purchased from Sigma- Aldrich.

**Cell metabolomics using QTRAP 5500**

Targeted metabolomics method, was used to quantitate >450 endogenous metabolites using QTRAP® 7500 LC-MS/MS System (Sciex, MA, USA). For the purpose, 100 μL of extraction buffer (methanol/water 50/50) containing 200 ng/mL of debrisoquine as internal standard for positive mode and 200 ng/mL of 4-nitrobenzoic acid as internal standard for negative mode was added to the cell pellet and sample tube was plunged into dry ice for 30 sec and 37 °C water bath for 90 sec. This cycle was repeated for two more times and then samples were sonicated for 1 minute. The samples were vortexed for 1 min and kept on ice for 20 minutes followed by addition of 100 μL of ACN. The samples were incubated at -20 °C for 20 minutes for protein precipitation. The samples were centrifuged at 13,000 rpm for 20 minutes at 4 °C. The supernatant was transferred to MS vial for LC-MS analysis. 20 μL of each prepared sample was mixed to generate the pooled QC sample.

**NIST Plasma sample preparation:** 25 µL of NIST plasma sample was dissolved in 100 μL of extraction buffer (methanol/water 50/50) containing 200 ng/mL of debrisoquine as internal standard for positive mode and 200 ng/mL of 4-nitrobenzoic acid as internal standard for negative mode. The sample was vortexed for 30 seconds and incubated on ice for 20 min followed by addition of 100 μL of acetonitrile and incubation at -20 ℃ for 20 min. Samples were centrifuged at 13,000 rpm for 20 min at 4 ℃. The supernatant was transferred to MS vial for LC-MS analysis.

One microliter of the prepared sample was injected onto a Kinetex F5, 2.6 μm 100 Å 150 × 2.1 mm (Phenomenex, CA, USA) using SIL-30 AC auto sampler (Shimazdu) connected with a high flow LC-30AD solvent delivery unit (Shimazdu) and CBM-20A communication bus module (Shimazdu) online with QTRAP 7500 (Sciex, MA, USA) operating in negative ion mode. A binary solvent comprising of water with 0.1% formic acid (solvent A) and acetonitrile with 0.1% formic acid (solvent B) was used. The extracted metabolite was resolved at 0.2 mL/min flow rate. The LC gradient conditions were as follows: Initial – 100% A, 0% B for 2.1 minutes; 14 minutes – 5% A, 95% B till 15 minutes; 15.1 minutes – 100% A, 0% B till 20 minutes. The auto sampler and oven were kept at 15 °C and 30 °C, respectively. Source and gas setting for the method were as follow: curtain gas = 40, CAD gas = 9, ion spray voltage = 1700 V in positive mode and ion spray voltage = 1600 V in negative mode, temperature = 350 °C, ion source gas 1 = 30 and ion source gas 2 = 50.

**Data Processing**

The data were normalized to respective internal standard area and processed using MultiQuant 3.0.3 (Sciex). The quality and reproducibility of LC-MS data was ensured using a number of measures. The column was conditioned using the pooled QC samples initially and were also injected periodically to monitor shifts in signal intensities and retention time as measures of reproducibility and data quality of the LC-MS data. We also ran NIST plasma sample, periodically, prepared in the same manner to check the instrumental variance. We also have blank solvent runs between set of samples to minimize carry-over effects. The report of pooled QC and NIST plasma is provided in excel sheet attached.
