## Supplemental Figures for "MAT2a and AHCY inhibition disrupts antioxidant metabolism and reduces glioblastoma cell survival"

### Slide 1
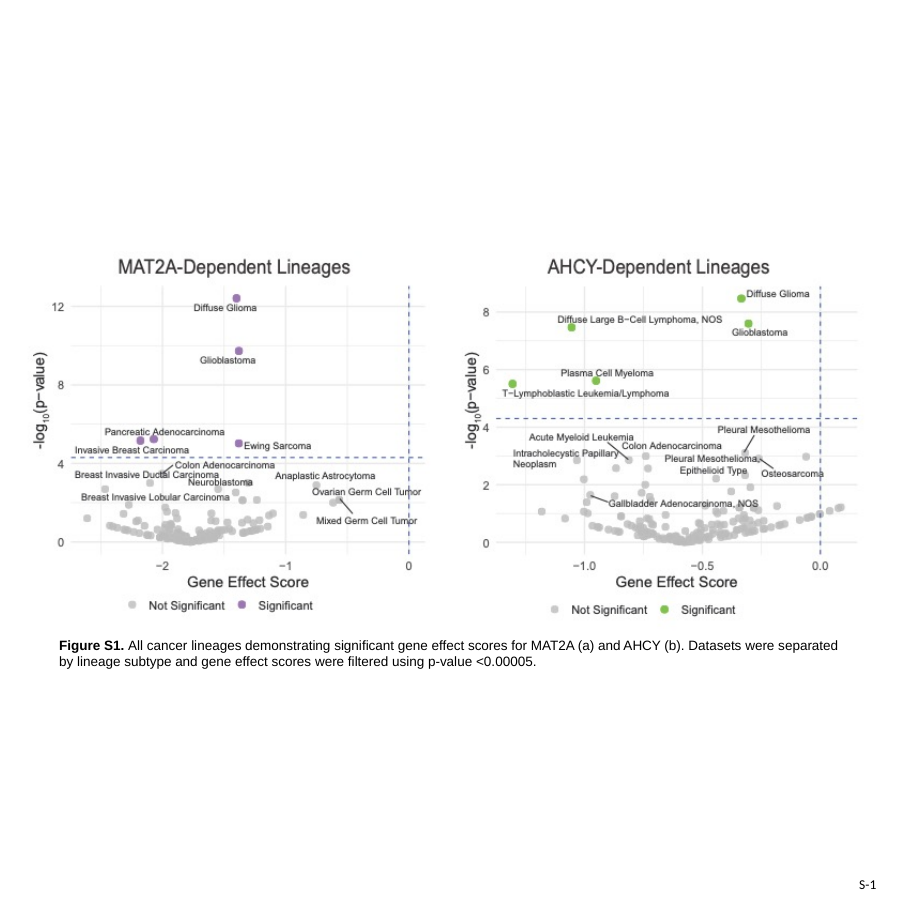

a
b
Figure S1. All cancer lineages demonstrating significant gene effect scores for MAT2A (a) and AHCY (b). Datasets were separated by lineage subtype and gene effect scores were filtered using p-value <0.00005.
S-1

### Slide 2
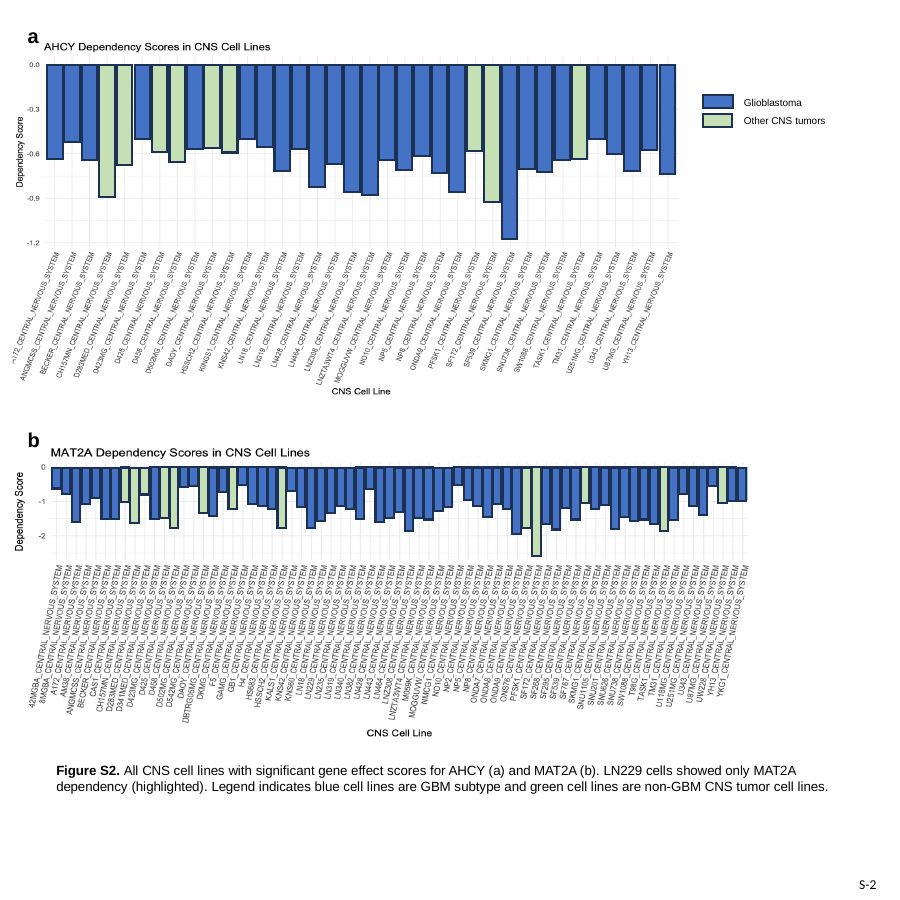

a
Glioblastoma
Other CNS tumors
b
Figure S2. All CNS cell lines with significant gene effect scores for AHCY (a) and MAT2A (b). LN229 cells showed only MAT2A dependency (highlighted). Legend indicates blue cell lines are GBM subtype and green cell lines are non-GBM CNS tumor cell lines.
S-2

### Slide 3
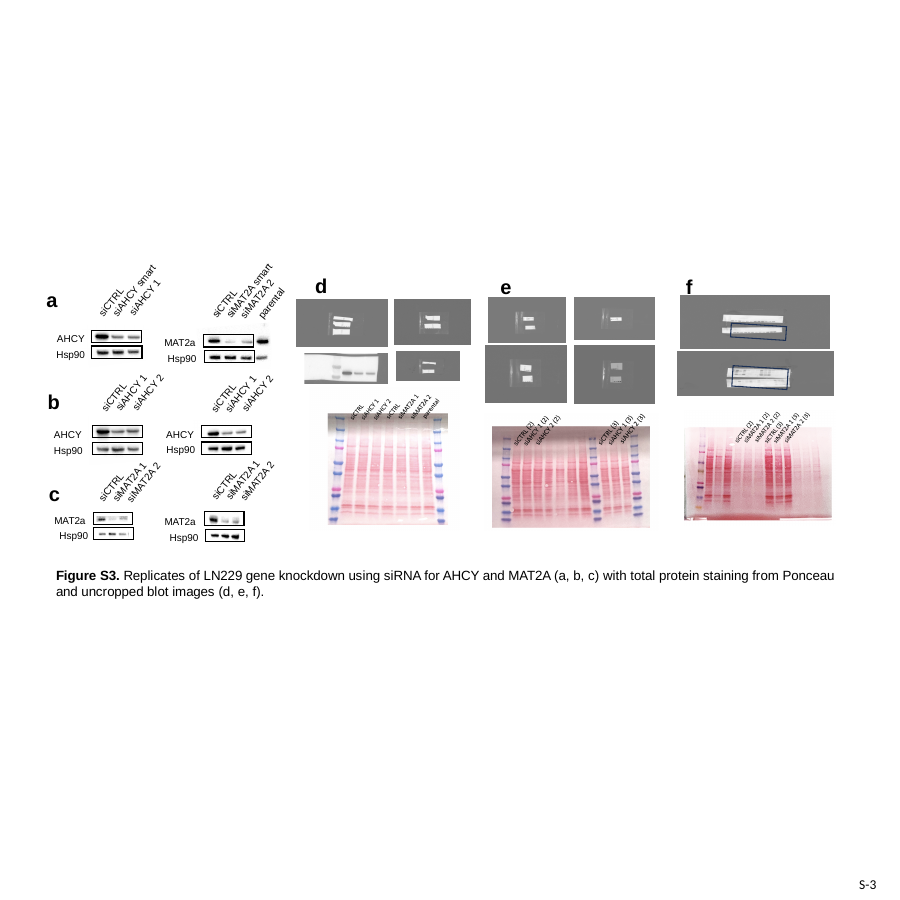

d
f
e
siAHCY 1
siAHCY smart
siCTRL
siMAT2A smart
siMAT2A 2
a
siCTRL
parental
AHCY
MAT2a
Hsp90
Hsp90
siCTRL
siAHCY 1
siAHCY 2
siCTRL
siAHCY 2
siAHCY 1
b
AHCY
AHCY
Hsp90
Hsp90
siMAT2A 1
siMAT2A 2
siCTRL
siCTRL
siMAT2A 1
siMAT2A 2
c
MAT2a
MAT2a
Hsp90
Hsp90
Figure S3. Replicates of LN229 gene knockdown using siRNA for AHCY and MAT2A (a, b, c) with total protein staining from Ponceau and uncropped blot images (d, e, f).
S-3

### Slide 4
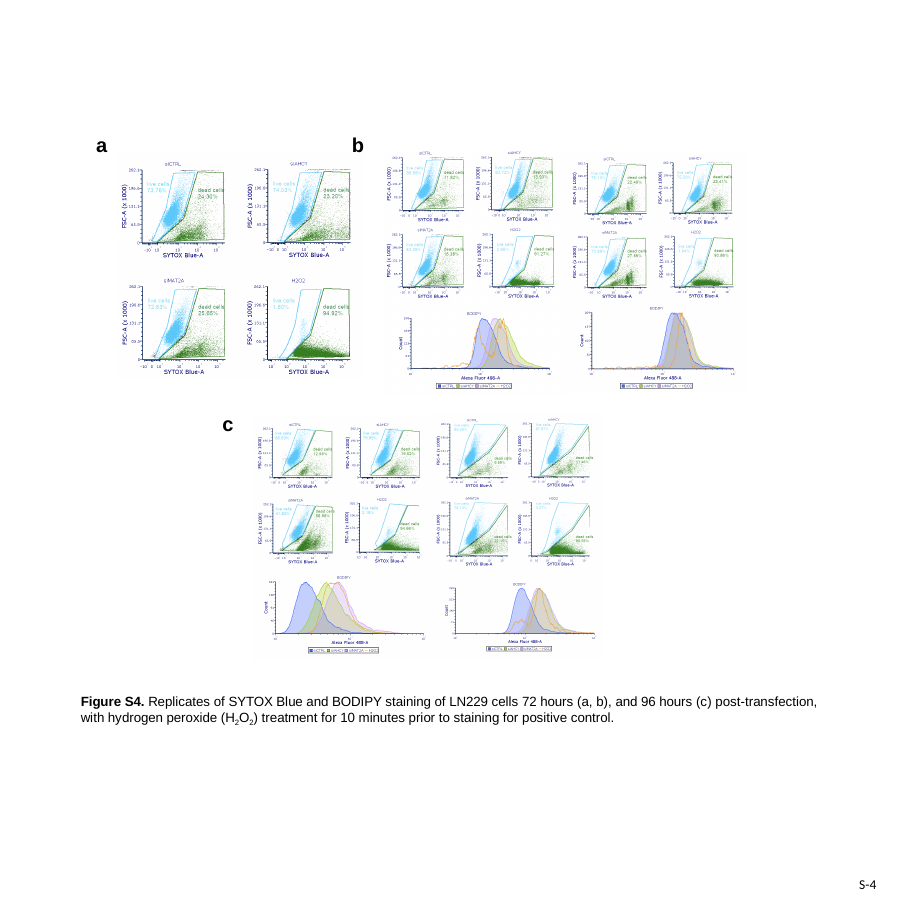

a
b
c
Figure S4. Replicates of SYTOX Blue and BODIPY staining of LN229 cells 72 hours (a, b), and 96 hours (c) post-transfection, with hydrogen peroxide (H_2_O_2_) treatment for 10 minutes prior to staining for positive control.
S-4

### Slide 5
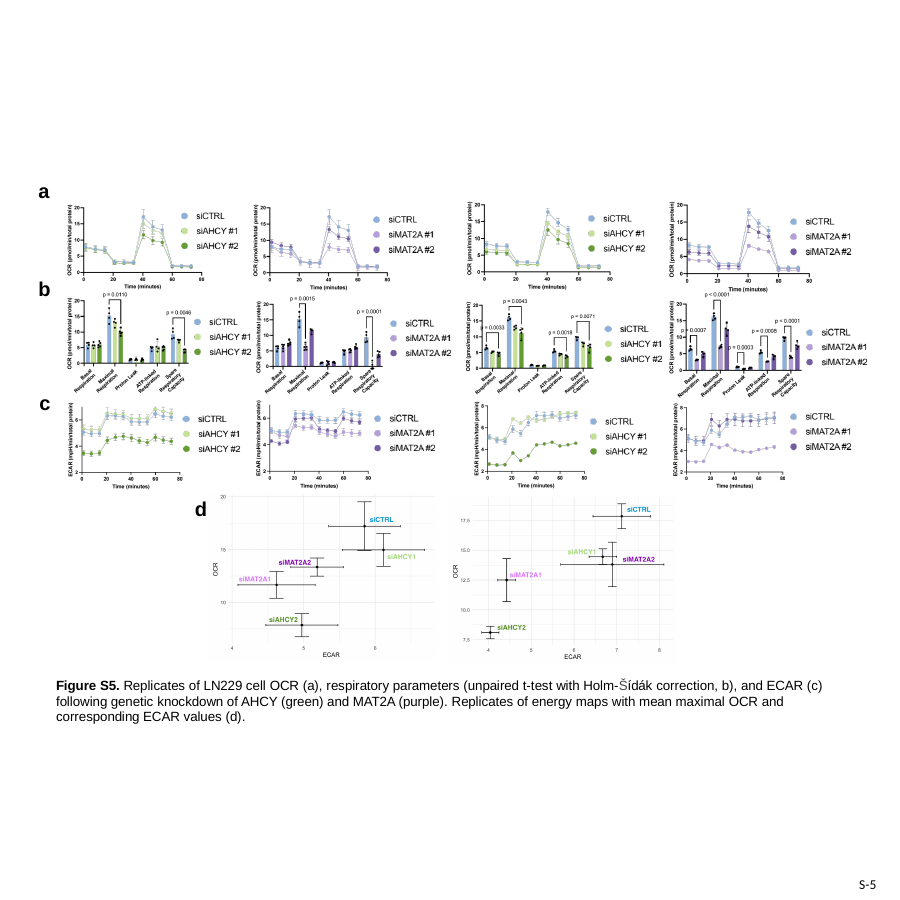

a
b
c
d
Figure S5. Replicates of LN229 cell OCR (a), respiratory parameters (unpaired t-test with Holm-Šídák correction, b), and ECAR (c) following genetic knockdown of AHCY (green) and MAT2A (purple). Replicates of energy maps with mean maximal OCR and corresponding ECAR values (d).
S-5

### Slide 6
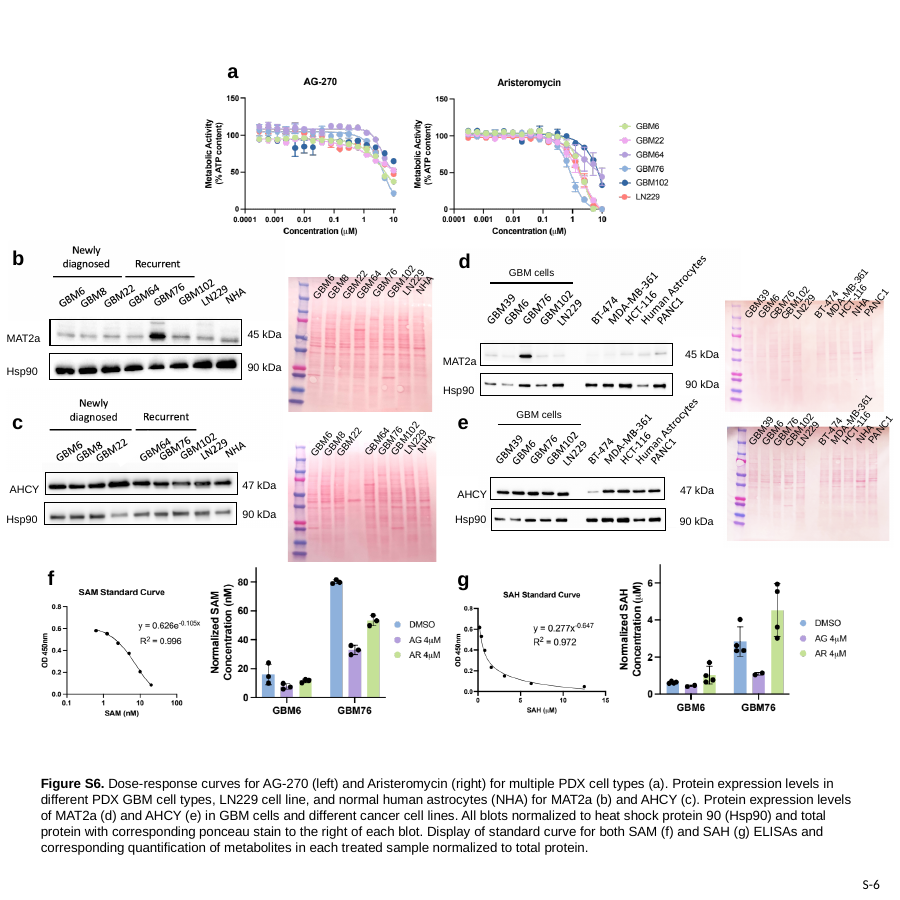

a
b
d
GBM cells
GBM102
GBM76
GBM64
LN229
GBM22
NHA
MDA-MB-361
GBM8
GBM6
GBM39
HCT-116
GBM102
PANC1
BT-474
GBM76
LN229
GBM6
NHA
45 kDa
MAT2a
45 kDa
MAT2a
90 kDa
Hsp90
90 kDa
Hsp90
GBM cells
MDA-MB-361
c
e
GBM39
HCT-116
GBM102
PANC1
BT-474
GBM76
LN229
GBM6
GBM6
NHA
GBM102
GBM76
LN229
GBM64
GBM22
NHA
GBM8
47 kDa
AHCY
47 kDa
AHCY
90 kDa
Hsp90
Hsp90
90 kDa
f
g
Figure S6. Dose-response curves for AG-270 (left) and Aristeromycin (right) for multiple PDX cell types (a). Protein expression levels in different PDX GBM cell types, LN229 cell line, and normal human astrocytes (NHA) for MAT2a (b) and AHCY (c). Protein expression levels of MAT2a (d) and AHCY (e) in GBM cells and different cancer cell lines. All blots normalized to heat shock protein 90 (Hsp90) and total protein with corresponding ponceau stain to the right of each blot. Display of standard curve for both SAM (f) and SAH (g) ELISAs and corresponding quantification of metabolites in each treated sample normalized to total protein.
S-6

### Slide 7
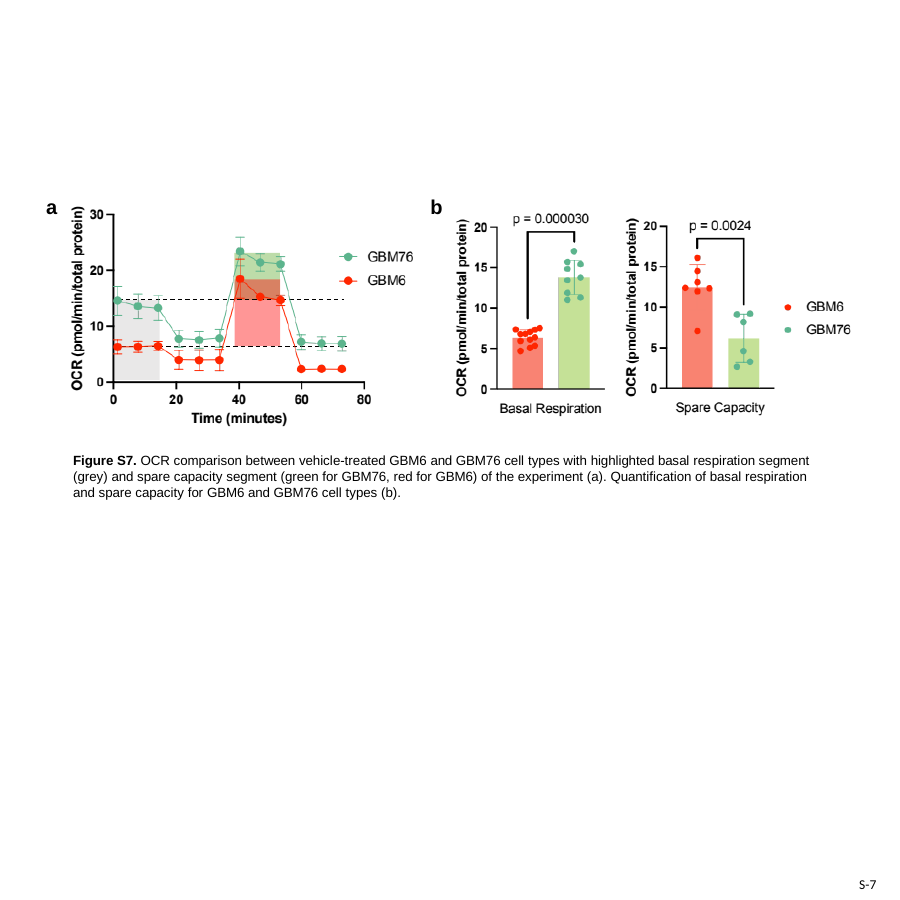

a
b
Figure S7. OCR comparison between vehicle-treated GBM6 and GBM76 cell types with highlighted basal respiration segment (grey) and spare capacity segment (green for GBM76, red for GBM6) of the experiment (a). Quantification of basal respiration and spare capacity for GBM6 and GBM76 cell types (b).
S-7

### Slide 8
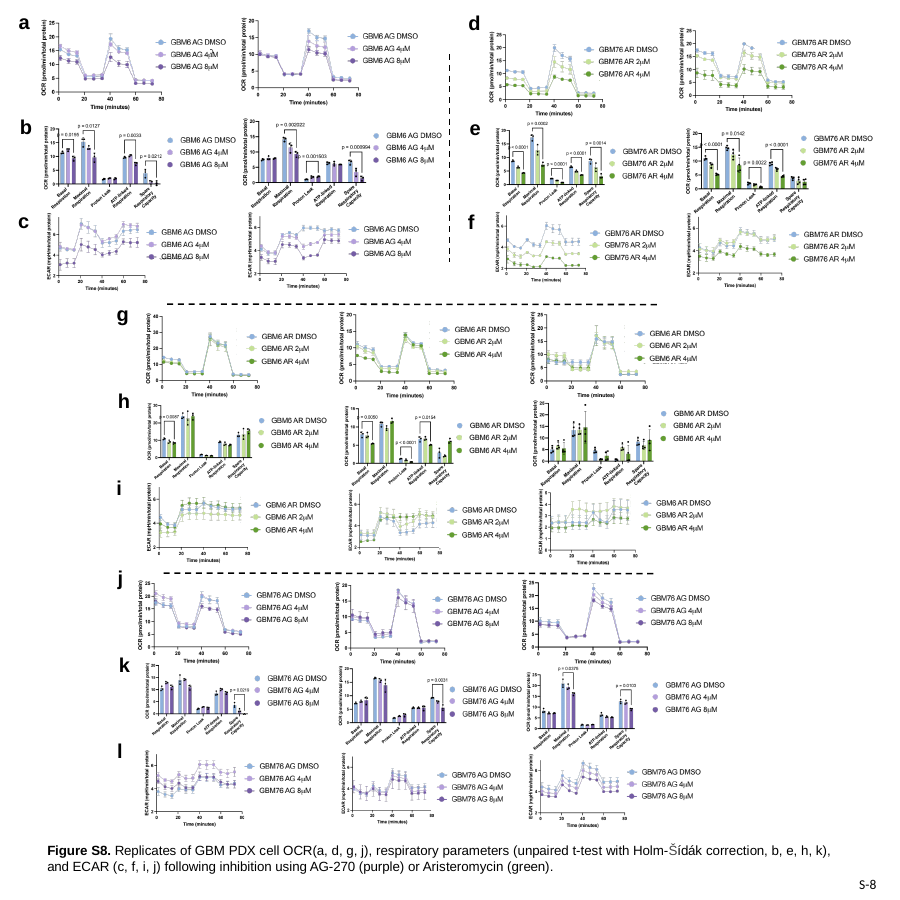

a
d
b
e
c
f
g
h
i
j
k
l
Figure S8. Replicates of GBM PDX cell OCR(a, d, g, j), respiratory parameters (unpaired t-test with Holm-Šídák correction, b, e, h, k), and ECAR (c, f, i, j) following inhibition using AG-270 (purple) or Aristeromycin (green).
S-8

### Slide 9
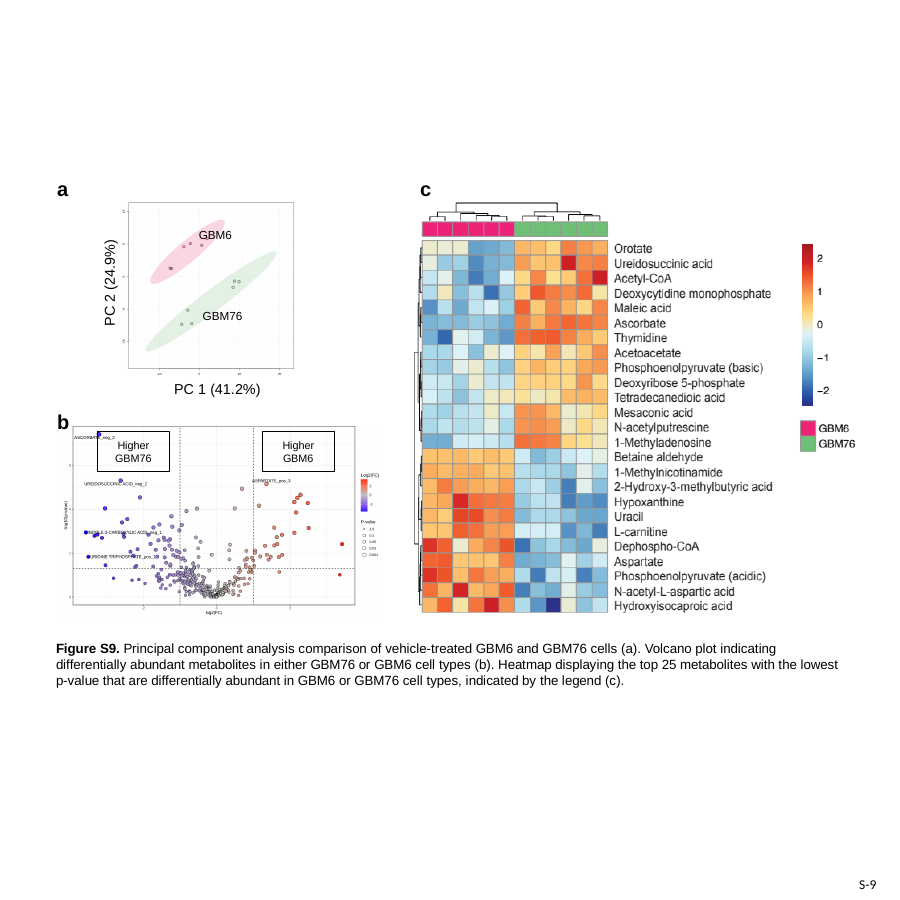

a
c
GBM6
PC 2 (24.9%)
GBM76
PC 1 (41.2%)
b
Higher
GBM6
Higher
GBM76
Figure S9. Principal component analysis comparison of vehicle-treated GBM6 and GBM76 cells (a). Volcano plot indicating differentially abundant metabolites in either GBM76 or GBM6 cell types (b). Heatmap displaying the top 25 metabolites with the lowest p-value that are differentially abundant in GBM6 or GBM76 cell types, indicated by the legend (c).
S-9

### Slide 10
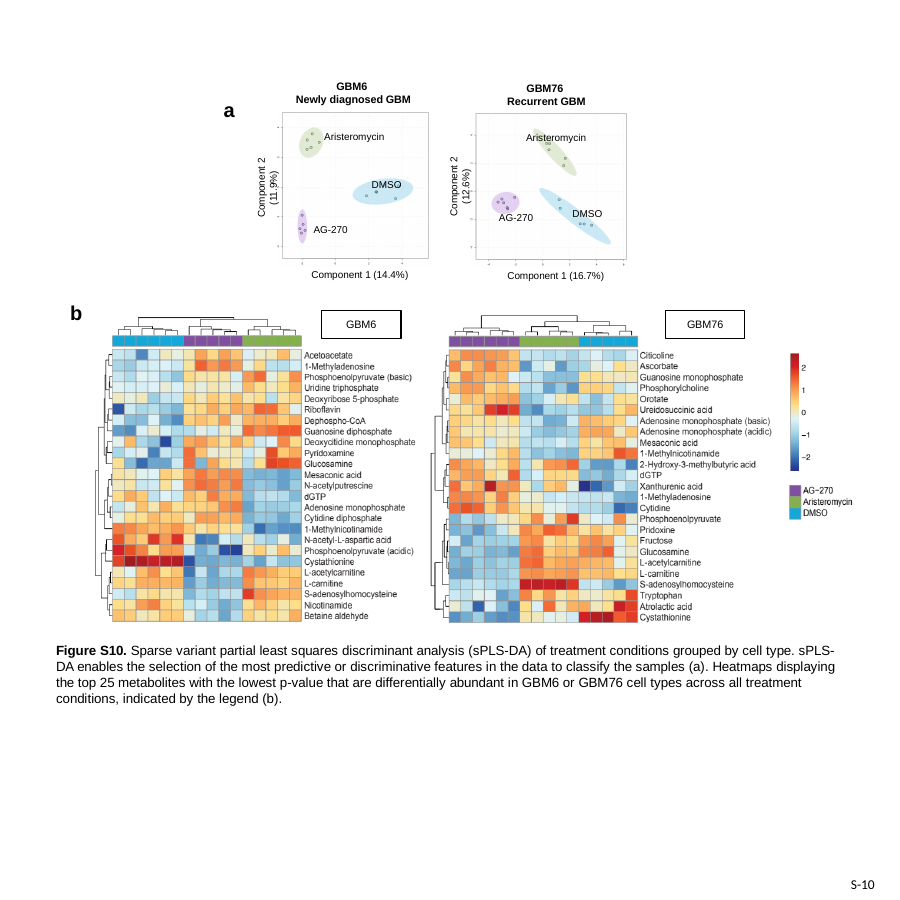

GBM6
Newly diagnosed GBM
Aristeromycin
Component 2 (11.9%)
DMSO
AG-270
Component 1 (14.4%)
GBM76
Recurrent GBM
Aristeromycin
Component 2 (12.6%)
DMSO
AG-270
Component 1 (16.7%)
a
b
GBM6
GBM76
Figure S10. Sparse variant partial least squares discriminant analysis (sPLS-DA) of treatment conditions grouped by cell type. sPLS-DA enables the selection of the most predictive or discriminative features in the data to classify the samples (a). Heatmaps displaying the top 25 metabolites with the lowest p-value that are differentially abundant in GBM6 or GBM76 cell types across all treatment conditions, indicated by the legend (b).
S-10

### Slide 11
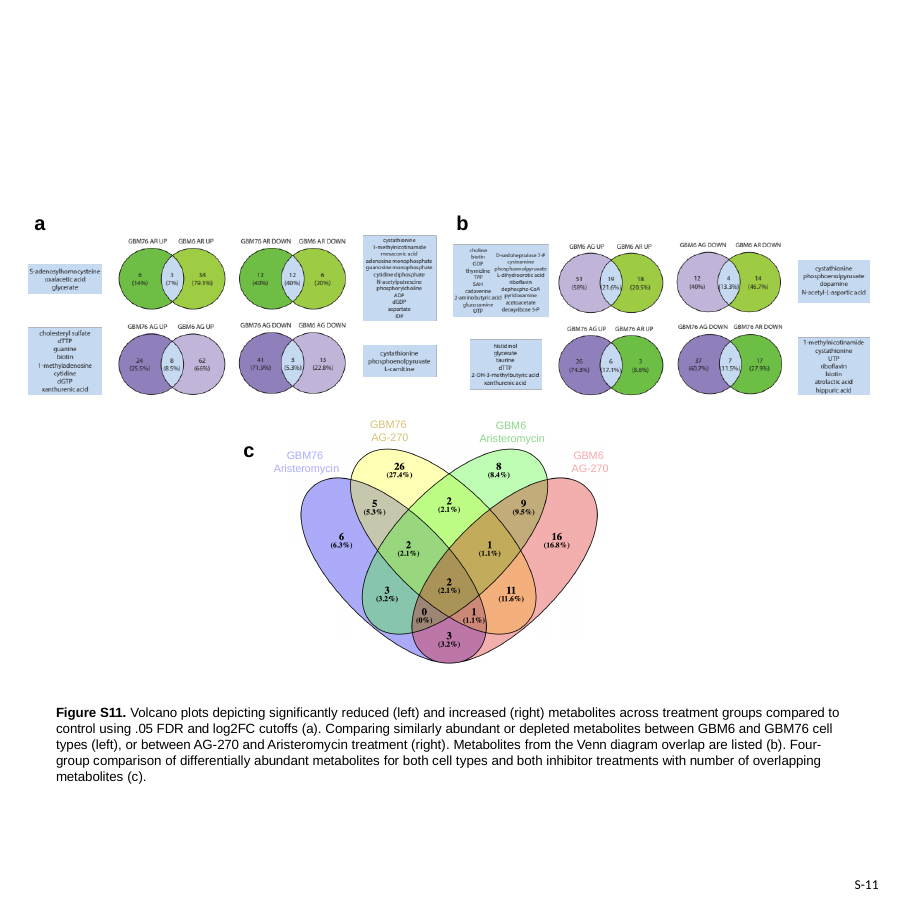

a
b
GBM76
AG-270
GBM6
Aristeromycin
GBM6
AG-270
GBM76
Aristeromycin
c
Figure S11. Volcano plots depicting significantly reduced (left) and increased (right) metabolites across treatment groups compared to control using .05 FDR and log2FC cutoffs (a). Comparing similarly abundant or depleted metabolites between GBM6 and GBM76 cell types (left), or between AG-270 and Aristeromycin treatment (right). Metabolites from the Venn diagram overlap are listed (b). Four-group comparison of differentially abundant metabolites for both cell types and both inhibitor treatments with number of overlapping metabolites (c).
S-11
